# BCL2L13 attenuation links impaired mitophagy to epithelial plasticity and anoikis tolerance in lung adenocarcinoma

**DOI:** 10.64898/2026.08.28.747809

**Authors:** Javad Alizadeh, Simone C. da Silva Rosa, Abhay Srivastava, Mahmoud Aghaei, Zeinab Babaei, Aleksandra Glogowska, Amir Barzegar-Behrooz, Amir Ravandi, Sabine Hombach-Klonisch, Sanjiv Dhingra, Michael Mowat, Rui Vitorino, Joseph W Gordon, Biniam Kidane, Naseer Ahmed, Saeid Ghavami

## Abstract

BCL2L13 is a mitochondrial BCL2-family protein linked to mitophagy and ceramide metabolism, but its role in NSCLC metastatic plasticity remains unclear. Human lung cancer Tissue Microarray (TMA) and matched patient specimens showed subtype- and site dependent BCL2L13 expression, with higher cytoplasmic/granular staining in primary NSCLC and reduced, heterogeneous staining in lymph-node metastases, most evident in adenocarcinoma and squamous-cell carcinoma. Because Epithelial-mesenchymal transition (EMT) and anoikis resistance are central requirements for metastatic dissemination, this primary to node attenuation provided the rationale to test BCL2L13 knockdown and overexpression in metastasis-relevant NSCLC models. In A549 and LLC cell liness, TGF beta 1 induced coordinated mitophagy and EMT with mitochondrial enrichment of BCL2L13. BCL2L13 knockdown impaired TGF beta 1 and carbonyl cyanide m chlorophenyl hydrazone (CCCP) associated mitophagy, reducing LC3 beta mitochondria colocalization, TOMM20 LAMP1 overlap and mitochondrial LC3 II/p62/TOMM20 turnover; BNIP3 and NIX redistribution did not compensate. BCL2L13 loss enhanced EMT-marker switching and migration, whereas overexpression partially opposed these changes. During detachment, BCL2L13 knockdown reduced anoikis-associated apoptosis despite preserved mitochondrial recruitment of BAX/BAK/BNIP3/NIX, altered BID processing, non parallel caspase activity and shifted FAK phosphorylation. Pharmacological autophagy modulation did not reverse this anoikis phenotype. Lipidomics identified adhesion-state-dependent ceramide synthases CerS2/CerS6-linked sphingolipid remodeling: BCL2L13 knockdown increased C24 linked sphingolipid species in attached cells but reduced C16/C24 ceramide-related profiles during anoikis. These findings identify BCL2L13 downregulation as a metastasis-associated mitochondrial–lipid state that limits mitophagic quality control while favoring EMT and detachment survival in NSCLC adenocarcinoma.

## Introduction

Lung cancer remains the leading cause of cancer-related deaths worldwide, driven primarily by its aggressive nature and high metastatic potential, particularly in non-small cell lung cancer (NSCLC), which accounts for over 80% of cases ^1,2^. Metastasis, a complex process involving epithelial-to-mesenchymal transition (EMT) and resistance to anoikis, is a critical factor in poor prognosis and limited therapeutic outcomes ^3^. Anoikis, a form of detachment-induced apoptosis, is essential for preventing dysregulated cell survival and metastatic dissemination. However, resistance to anoikis enables cancer cells to invade distant tissues, contributing to metastasis ^4,5^.

Mitophagy, a selective autophagy process for clearing damaged mitochondria, is closely linked to cancer progression. Dysregulated mitophagy can impair mitochondrial function, promoting EMT and metastasis ^6,7^. Ceramides, synthesized by ceramide synthases (CerS1–6), play a pivotal role in regulating both mitophagy and apoptosis ^8^. Among ceramide synthases, CerS2 and CerS6 have been implicated in tumor migration and metastasis in NSCLC, with altered ceramide profiles driving cancer cell invasiveness ^9^. Despite the known roles of ceramides, the regulatory mechanisms connecting CerS activity to mitophagy and anoikis remain underexplored.

BCL2L13, a BCL2 family protein, is a key regulator of mitophagy and apoptosis and has emerged as a critical mediator in cancer progression. It has been shown to inhibit CerS2 and CerS6 activity by disrupting their heterodimerization, thereby reducing ceramide synthesis and promoting tumor growth in glioblastoma ^10^. However, the role of BCL2L13 in NSCLC metastasis and its regulation of ceramide-mediated mitophagy and anoikis has yet to be elucidated.

Despite significant advancements in understanding the molecular drivers of lung cancer, the mechanisms underlying metastasis, particularly in non-small cell lung cancer (NSCLC), remain incompletely understood, posing a major barrier to improving patient outcomes. While the critical roles of mitophagy and anoikis resistance in promoting metastasis have been established, the regulatory pathways that link these processes to ceramide metabolism are poorly defined. Specifically, ceramide synthases (CerS2 and CerS6) have been implicated in NSCLC progression, yet how their activity is modulated to drive tumor invasiveness and resistance to apoptosis remains largely unexplored. Moreover, while BCL2L13 has been recognized as a dual regulator of mitophagy and apoptosis, its ability to influence CerS2 and CerS6 activity, and its subsequent impact on metastatic phenotypes such as epithelial-to-mesenchymal transition (EMT) and anoikis resistance, remains uncharacterized in NSCLC. This gap in knowledge is critical, as understanding the interplay between BCL2L13, ceramide metabolism, and mitophagy could unveil novel therapeutic strategies targeting metastatic lung cancer. Addressing this gap, our study seeks to elucidate the role of BCL2L13 in modulating CerS2- and CerS6-dependent ceramide synthesis, mitophagy, and anoikis resistance in lung adenocarcinoma. By defining these mechanisms, we aim to provide insights into the molecular drivers of NSCLC metastasis and identify actionable targets for therapeutic intervention.

## Materials & Methods

### Reagents and Antibodies

Cell culture plastic ware (adherent and repellent), media, penicillin, streptomycin, fetal bovine serum (FBS), and Geneticin (G418 sulfate) were sourced from VWR (Toronto, ON, Canada). Bafilomycin A1 (B1793), Rapamycin (R8781), propidium iodide (PI), MTT, protease and phosphatase inhibitors, as well as anti-rabbit and anti-mouse peroxidase-conjugated secondary antibodies, were obtained from Sigma-Aldrich (Oakville, CA). Mitophagy inducer CCCP, TMRM, MitoTracker™ Red CMXRos, and MitoSOX™ Mitochondrial Superoxide Indicators were purchased from Thermo Fisher Scientific (Waltham, MA, USA). Caspase-Glo® 3/7, 8, and 9 assays were obtained from Promega (Toronto, ON, Canada). Recombinant human and mouse TGFβ1 (240-B, 5 ng/ml) were from R&D Systems (Mississauga, ON, Canada). The pcDNA3 plasmid was acquired from Addgene (Watertown, MA, USA), and the jetPRIME® DNA transfection reagent was provided by Dr. Joseph Gordon (University of Manitoba). BCL2L13/control shRNA, puromycin, and polybrene were purchased from Santa Cruz Biotechnology (Dallas, TX, USA), along with GAPDH and Lamin A/C monoclonal antibodies. BCL2L13 polyclonal antibody was from Proteintech (Rosemont, IL, USA), while LC3, BNIP3, and β-Actin antibodies were from Sigma-Aldrich. Other primary antibodies were sourced from Cell Signaling Technology (Canada), and AlexaFluor-488/594-conjugated secondary antibodies were purchased from Jackson Lab (CA, USA).

### Cell Culture

For all experiments we used the following lung carcinoma cell lines: lung adenocarcinoma (A549 (ATCC-CCL-185™), and mouse Lewis Lung Carcinoma (LL/2-LLC1- (ATCC® CRL1642). Cancer cells were cultured in Dulbecco’s Modified Eagle’s Medium high glucose (4 mg/ml) (DMEM) (CORNING; Cat #: 50-003-PB) supplemented with 10% Fetal Bovine Serum (FBS) (Gibco™; Cat #: 16000044) and 1% penicillin and streptomycin. Insulin/Transferrin/Selenium (ITS) (1%) (Gibco™; #41400045) was used to starve the cells and avoid the starvation induced autophagy. Cells were maintained in a humidified incubator with 95% air and 5% CO2 at 37 °C. Stable *BCL2L13* knockdown (KD) and *BCL2L13* overexpressing (OE) cells were grown in the same medium plus puromycin (4 μg/ml) or Geneticin (750 nM) for each cell type, respectively ^11^.

### Cell Viability Assay

We measured the viability of A549 and LLC cancer cells (scramble, *BCL2L13-*knockdown (KD), and *BCL2L13-*overexpressing (OE)) under treatment with various concentrations of CCCP using MTT assay, as described previously ^12–14^. Briefly, A549 and LLC cancer cells were treated with different concentrations of CCCP (0–20 μM, 48 hrs) and Bafilomycin A1 (0-10nM, 48 hrs). Relative cell viability (percent of control) was calculated using the equation: (mean OD (570nm) of treated cells/mean OD (570nm) of control cells) × 100. The treated cells were compared with control cells that had been treated with dimethyl sulfoxide (DMSO) vehicle only.

### Measurement of Apoptosis by Flow Cytometry

The Nicoletti method was used to measure cellular apoptosis ^15–17^. A549 and LLC cancer cells (control, *BCL2L13*-KD, *BCL2L13*-OE) were cultured in 12-well plates and treated with CCCP (0–20 μM) or Bafilomycin A1 (0–10 nM) for 48 hours. Cells were detached using EDTA buffer, harvested by centrifugation at 1500 g for 5 minutes at 4°C, washed in PBS, and resuspended in hypotonic PI lysis buffer (0.1% Triton X-100, 1% sodium citrate, 0.5 mg/ml RNase A, 40 μg/ml propidium iodide). After incubation at 37°C for 30 minutes, nuclei were analyzed using a Beckman Coulter Cytoflex LX flow cytometer (488 nm blue laser, 610/20 BandPass filter). Single-cell populations were gated, and cell cycle analysis quantified the percentage of cells in Sub-G0, G0-G1, S, and G2-M phases, identifying apoptotic nuclei as hypo-diploid DNA on the left of the G1 peak.

### Luminometric Caspase Assay

Caspase-Glo®-3/-7, -8, and -9 (Promega) was used for the determination of proteolytic activity of caspase-8, 9, 3/7 in luminometric assays according to the manufacturer’s instructions and our previous report ^13,18^. A549 and LLC cancer cells (control, *BCL2L13*-KD, *BCL2L13*-OE) were cultured in 96-well plates (15,000 cells/well) for 72 hours (caspase 8, 9) or 96 hours (caspase 3/7). Fresh caspase reagents (z-LETD-Luciferin, z-DEVD-Luciferin, z-LEHD-Luciferin) and protein cell lysate buffer were prepared, with medium-only and reagent blank controls included. After gentle shaking (300–500 rpm, 30 seconds) and 90 minutes of room temperature incubation, the solution was transferred to white-well plates, and luminescence was measured relative to controls.

### MitoSOX Staining for Measurement of Mitochondrial ROS

To assess mitochondrial ROS, A549 and LLC cancer cells (control, *BCL2L13*-KD, *BCL2L13*-OE) were cultured in 6-well plates for 24 hours and stained with MitoSOX™ Red (1.25 µM) and Hoechst nuclear stain (10 μM) at 37 °C for 30 minutes. After washing, cells were imaged using a fluorescence microscope, and fluorescence intensity was quantified in at least 50 individual cells per condition using ImageJ software (NIH, Bethesda, MD, USA).

### TMRM Staining for Measurement of Mitochondrial Membrane Potential

Mitochondrial membrane potential, essential for mitochondrial function, was assessed using the cell-permeant dye TMRM (100 nM), which accumulates in active mitochondria and fluoresces. A549 and LLC cancer cells (control, *BCL2L13*-KD, *BCL2L13*-OE) were cultured in 6-well plates for 24 hours, stained with TMRM and Hoechst (10 μM) at 37 °C for 30 minutes, and washed. Fluorescence images were captured, and intensity was quantified using ImageJ software (NIH, Bethesda, MD, USA) by averaging measurements from 50 randomly selected cells per condition.

### Immunoblotting

Western blotting was performed as described previously ^11,13^, with β-actin and GAPDH as loading controls. Protein extracts were prepared in lysis buffer, quantified via Lowry protein assay, separated by SDS-PAGE, and transferred to nylon membranes. Membranes were blocked, incubated overnight with primary antibodies at 4°C, followed by HRP-conjugated secondary antibodies for 2 hours at room temperature. Bands were visualized using ECL detection with a Bio-Rad imager, and signals were quantified using Alpha Ease FC software.

### Subcellular Fractionation

Protein samples were extracted using lysis buffer with protease and phosphatase inhibitors (Sigma-Aldrich, CA). Subcellular fractionation was performed using mitochondrial (Qiagen Qproteome) and nuclear/cytoplasmic extraction kits (Thermo Fisher). Proteins were quantified by Lowry assay and analyzed by immunoblotting. Primary antibodies (1:1000) and HRP-conjugated secondary antibodies (1:5000 Goat Anti-Rabbit; 1:3000 Goat Anti-Mouse) were used with chemiluminescence for visualization on a Bio-Rad imager. COX IV and Lamin A/C confirmed fractionation purity, and Image Lab Software ensured linear exposure.

### Production of Stable BCL2L13 Knockdown (KD) Cells

A549 and LLC cells (5 × 10⁴ cells/well) were seeded in 12-well plates and cultured in DMEM with 10% FBS for 24 hrs. At 40% confluency, cells were treated with 10 μg/ml polybrene (Santa Cruz Biotechnology) in serum-free DMEM for 1 hr, then transfected overnight with lentiviral particles encoding *BCL2L13* shRNA or scrambled control (puromycin-resistant). Transfection was performed at 12, 24, and 32 MOI, followed by 24 hrs recovery in fresh medium. Stable transfectants were selected with puromycin (4 μg/ml), and BCL2L13 expression was confirmed by Western blot in stable clones.

### Production of BCL2L13 Overexpressing (OE) Cells

A549 and LLC cells (5 × 10⁴ cells/well) were seeded in 12-well plates and cultured in DMEM with 10% FBS for 24 hrs. At 40% confluency, cells were transfected with 0.8 μg of pcDNA3 encoding *BCL2L13* using JetPrime Polyplus reagent (Polyplus, NY, USA) per manufacturer’s instructions. After 24 hrs, the medium was replenished, and cells recovered for 48 hrs before selection in Geneticin (750 nM). BCL2L13 expression was confirmed by Western blot in five stable clones, with the highest-expressing clones used for subsequent experiments.

### Immunocytochemistry (ICC)

To assess autophagy/mitophagy changes in A549 and LLC cells, including *BCL2L13* knockdown transfectants, immunocytochemistry (ICC) was performed as previously described ^19,20^. Cells (5000/spot) were cultured on coverslips in DMEM with 10% FBS, starved with 1% ITS for 24 hrs, and treated with TGFβ1 (5 ng/ml) for 36 hrs. At the indicated time, cells were stained with MitoTracker Red (200 nM) and Hoechst (10 μM) for 30 mins at 37°C, fixed with 3.7% formaldehyde, permeabilized with 0.25% Triton-X100, and blocked with 5% normal donkey serum. Overnight incubation at 4°C was done using rabbit anti-LC3, anti-TOMM20, or anti-LAMP1 primary antibodies and isotype controls (Cell Signaling). After washing, cells were incubated with Alexa Fluor-conjugated secondary antibodies (488 and 594), mounted with nuclear counterstain, and stored at -20°C. Imaging and quantification were performed using ImageJ and Zen 2.3 Pro software.

### Tissue Microarray (TMA)

A lung tissue array kit (NBP2-30222 lot 4; Novus Biologicals, Canada) containing lung samples (primary lung tumor, metastatic tumor, and healthy lung) was purchased from US Biomax, Inc. Arrays contained different types of lung cancer including squamous cell carcinoma, lung adenocarcinoma, bronchioloalveolar carcinoma, adenosquamous carcinoma, large cell carcinoma, and small cell carcinoma). The information on all tissue samples, clinical stages and pathology grades were provided. Slides were processed by performing IHC for BCL2L13 (Protein Tech, 16612-1-AP, 1:100 dilution) followed by Tissue Microarray Scoring as previously described ^21^. IHC results were blindly evaluated by three independent pathologists who scored the samples based on the intensity of the staining [none (N), weak (W), moderate (M), and strong (S)].

#### Human Study design, ethics, participant recruitment, and tissue selection

This translational tissue study was conducted within the University of Manitoba clinical protocol entitled “The Potential Application of Mitophagy Signature as a Novel Prognostic Approach in Lung Cancer Metastasis.” The protocol received final approval from the University of Manitoba Biomedical Research Ethics Board on 9 October 2018 (BREB No. HS21943 [B2018:066]). All participants provided written informed consent before enrolment. The parent clinical cohort was recruited between 2019 and 2023 through CancerCare Manitoba and the Thoracic Clinical Program at Health Sciences Centre, Winnipeg, Manitoba, Canada.

For the present analysis, eight patient-matched pairs of primary lung tumour and regional lymph-node tissue were selected from the clinical cohort. Each pair comprised the primary lung tumour, designated P, and the corresponding lymph node from the same patient, designated N. All lymph-node specimens contained histologically confirmed metastatic tumour. The paired cohort included adenocarcinoma, squamous cell carcinoma, and small-cell carcinoma. Non-neoplastic lung and lymph-node tissues were additionally examined as reference controls where available. Histological classification and the presence of metastatic tumour in the lymph nodes were confirmed using the corresponding diagnostic pathology assessment before BCL2L13 analysis.

#### BCL2L13 immunohistochemistry in matched primary tumour–lymph-node specimens

BCL2L13 protein expression was evaluated by immunohistochemistry using an anti-BCL2L13 primary antibody from Proteintech at a 1:100 dilution. Tissue sections were processed according to the laboratory’s validated immunohistochemistry protocol, and antibody binding was visualized using a brown 3,3′-diaminobenzidine chromogen with hematoxylin nuclear counterstaining. Primary tumours and their matched metastatic lymph nodes were evaluated comparatively within each patient pair.

BCL2L13 positivity was defined as cytoplasmic or granular brown staining within morphologically identifiable tumour cells. Staining intensity was categorized as absent, weak, moderate, or strong. Non-neoplastic lung tissue was assessed for basal epithelial and stromal staining, whereas non-neoplastic lymph-node tissue served as a reference for background and non-tumour-associated signal. Pigment, necrotic or keratinized material, macrophage-associated staining, and isolated nonspecific punctate deposits that did not correspond morphologically to viable tumour cells were excluded from interpretation. The patient pair, rather than an individual microscopic field, was treated as the biological unit of analysis.

### Transmission Electron Microscopy (TEM)

Autophagy and mitophagy induction were confirmed using TEM based on a protocol described previously ^12^. Briefly, A549 and LLC cells were cultured in 100 mm dishes (250,000 cells/dish) in DMEM media (high glucose) with 10% FBS in standard cell culture incubator conditions. Cells were starved with ITS (1%) for 24 h and then treated with TGFβ1 (5 ng/ml) for 36 h. TEM was performed on ultra-thin sections (100 nm on 200 mesh grids) 36 hrs after treatment and sections were stained with uranyl acetate and counterstained with lead citrate. Autophagy or mitophagy induction was evaluated based on the autophagosome and autophagolysosome formation in TEM images.

### Measurement of TGFβ1 Concentration in Cell Culture Media

*BCL2L13*-WT A549 and *Bcl2l13*-WT LLC cells were cultured in 100 mm dishes (250,000 cells/dish) in DMEM media (high glucose) with 10% FBS in standard cell culture incubator conditions at 37 °C and 5% CO2 for 24 hrs. Supernatants were collected and then concentrated using the Amicon Ultra-4 centrifugal filters (3000 NMWL) for 1 hr at 4 °C. Collected concentrates were then frozen at -80 °C until shipped to Eve Technologies (Calgary, AB, Canada) for assessment of human TGFβ1 and mouse TGFβ1, using the “TGFB-Plex Discovery Assay® Multi Species Array”. Data were statistically analyzed using GraphPad Prism.

### Cell Invasion and Migration (CIM) Assay

The xCelligence RTCA DP system (ACEA Biosciences, Inc.) was used to monitor real-time migration of A549 and LLC cells (scramble vs. *BCL2L13*-KD). This system employs microelectronic sensors on the underside of a Boyden-like chamber membrane to detect cell migration. As cells migrate from the upper to the lower chamber, impedance changes are recorded, reflecting the number of migrated cells. A549 and LLC cells were cultured to 30–40% confluency in DMEM (10% FBS), starved for 24 hrs, and seeded (2×10⁴ cells/well) in the upper chamber, with 10% FBS in the lower chamber. Migration was tracked by the DP analyzer every minute for 24 hrs.

### Measurement of Mitochondrial Respiration

Mitochondrial respiration was measured using the Agilent Seahorse XFe24 analyzer by recording the Oxygen consumption rate (OCR). Approximately 0.3×10^6^ cells (control, *BCL2L13*-KD, and *BCL2L13*-OE) per well were seeded in their growth medium in a XF cell culture 24 well microplates. On the day of the experiment, cells were washed twice with and changed to XF base minimal DMEM media containing glucose, L-glutamine and sodium pyruvate (Agilent Technologies, Mississauga, ON). The cells were incubated at 37°C for 1 hr prior to starting the assay. Seahorse analyzer measured the baseline OCR first, followed by proton leak using 1 µM of oligomycin. Maximal respiration was determined by injection of FCCP (2 µM) and nonmitochondrial respiration by addition of 1 µM rotenone and 1 µM antimycin A together. Maximal respiration was calculated by subtracting non-mitochondrial OCR from FCCP OCR. Spare capacity was determined by subtracting baseline OCR from FCCP OCR. Further, ATP production was calculated by subtracting Oligomycin OCR from baseline OCR. Finally, proton leak was measured by subtracting non-mitochondrial OCR from Oligomycin OCR. Data are presented as pmol of oxygen/minute/cell count.

### Liquid Chromatography-Mass Spectrometry (LC-MS) for Lipidomics Analysis of Ceramide Species

Lipids were extracted using chloroform:methanol (2:1, v/v) from A549 and LLC cells (control, *BCL2L13*- KD,-OE) grown to 100% confluency. Cell homogenates were mixed with internal standards, centrifuged, and the lower phase dried under N₂ gas. Lipids were reconstituted in water-saturated butanol, sonicated, and mixed with methanol containing ammonium formate. Samples were prepared for LC-MS analysis, with separation on a Zorbax C18 column using a gradient of mobile phases A and B, both containing 10 mM ammonium formate. Lipids were analyzed on an AbSciex 4000 QTRAP mass spectrometer in positive ESI mode using MRM to screen 322 lipids across 25 classes. Lipids were identified by class-specific precursor ions or neutral losses and quantified against internal standards. Chromatographic and mass spectrometry parameters ensured precise separation of lipid species, including isobaric and molecular variants ^22–24^.

### Caspase Glu Luminometric Assay

Caspase-Glo-9, -3/7, and -8 (Promega) were used to measure the proteolytic activity of caspase-3/7, -9, and -8 as previously described ^25^.

### Statistical Analysis

Data were collected from three replicates across three independent experiments, with error bars representing ± standard deviation. Statistical analysis included one-way or two-way ANOVA with Tukey’s test for multiple comparisons. Kaplan-Meier survival analysis was performed using cBioPortal, stratifying tumors by *BCL2L13* mRNA expression, with data from TCGA and Legacy Firehose datasets. Correlation studies employed Kendall, Mann-Whitney, and ordinal regression tests.

#### Lipidomics Analysis

Lipid classification was performed using the LIPIDMAPS database, and data were analyzed via MetaboAnalyst 5.0. Hierarchical clustering heatmaps displayed lipid abundance, while PCA and PLSDA identified differential lipids between primary and metastatic groups (VIP>1.2, FDR<0.05). Swiss Target Prediction linked key lipids to interacting proteins, with functional annotation conducted via GO, KEGG, and protein-protein interaction analysis using String and GProfiler. Statistical significance was determined using a two-tailed t-test or ANOVA, with p<0.05 considered significant ^24^.

## Results

### TGFβ1 Drives Coupled Mitophagy–EMT Programs in NSCLC via BCL2L13-Linked Mitochondrial Remodeling

TGFβ1, a key inducer of EMT and metastasis in NSCLC ^11^, operates in an autophagy-dependent manner, where inhibition shifts cells toward epithelial phenotypes ^11^. Consistent with this, we previously observed mitochondria-like structures within cytosolic double membranes in TGFβ1-treated NSCLC cells^11^. Given the established role of mitophagy in metastasis ^26,27^, we assessed whether TGFβ1 coordinately induces mitophagy and EMT in A549 and LLC models.

TGFβ1 robustly increased early and late mitophagy, evidenced by LC3β–MitoTracker and TOMM20–LAMP1 co-localization (Figure 1A–D; Suppl. Figure 1A–D), alongside mitochondrial LC3β-II accumulation, p62 degradation, and TOMM20 loss in both A549 and LLC model (Figure 1E–H; Suppl. Figure 1E–H). Mechanistically, TGFβ1 enhanced mitochondrial localization of BCL2L13 (Figure 1I; Suppl. Figure 1I), while NIX and BNIP3 responses were cell-line specific (Figure 1J–M; Suppl. Figure 1J–M). Endogenous TGFβ1 secretion was confirmed in both models (Figure 1N; Suppl. Figure 1N). Concurrently, EMT was induced, marked by decreased E-cadherin and increased vimentin and N-cadherin (Figure 1O–R; Suppl. Figure 1O–R). Collectively, these results demonstrate that TGFβ1 induces both mitophagy and EMT in NSCLC cells, with increased mitochondrial BCL2L13 localization, suggesting its potential relevance to NSCLC metastasis.

**Figure 1.**
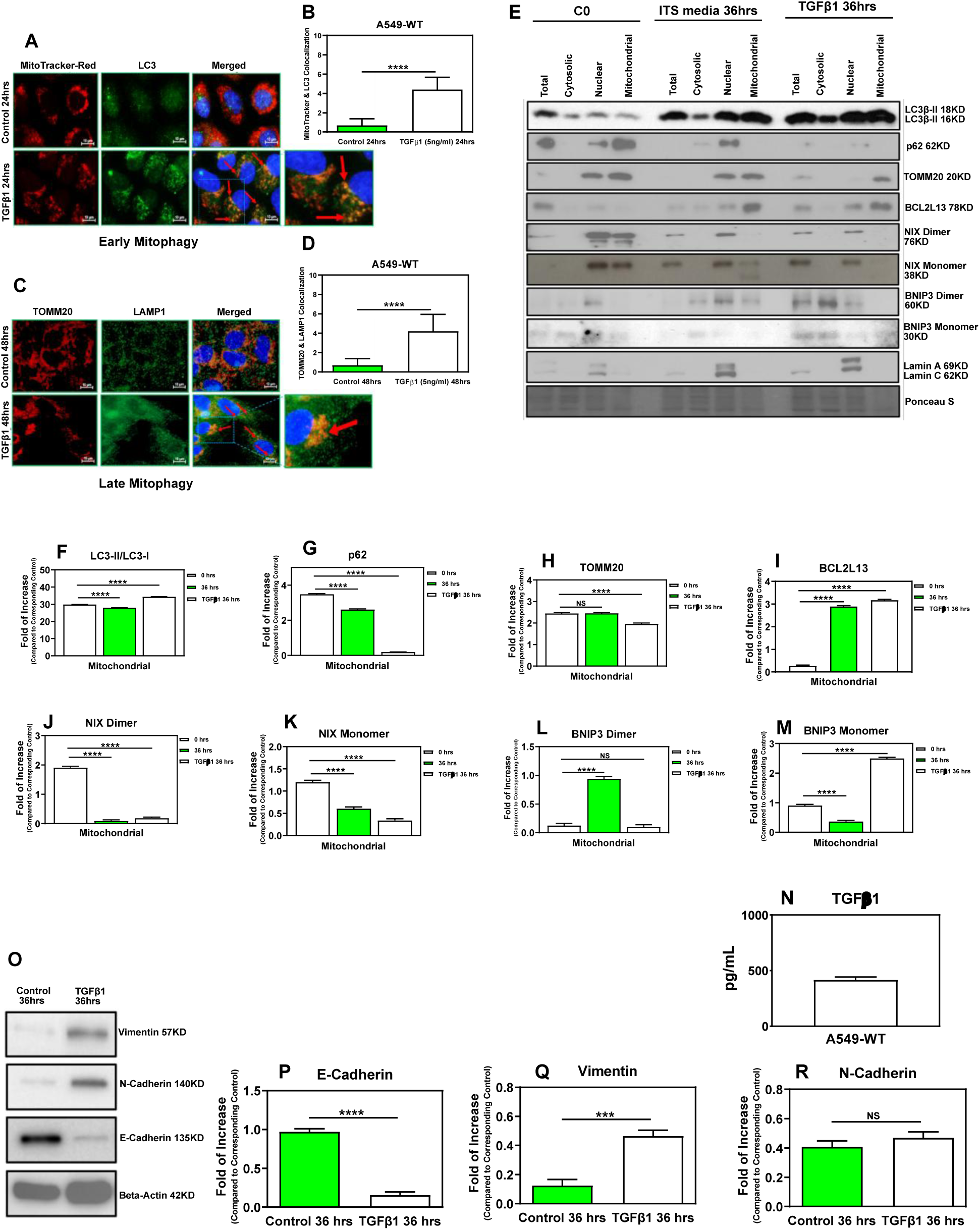
TGFβ1 Induces Coordinated Mitophagy and EMT in A549 Cells. 549 cells were treated with TGFβ1 (5 ng/ml) for 24 h (early mitophagy) or 48 h (late mitophagy) and analyzed by ICC using MitoTracker/LC3β (A) or TOMM20/LAMP1 (C) (scale bars: 10 μm), with quantification of co-localization shown in (B, D) (n=10 fields, ∼5 cells/field). Subcellular fractionation followed by immunoblotting at 36 h (E) assessed mitophagy markers, with Lamin A/C and Ponceau S confirming fraction purity and loading, and mitochondrial protein levels quantified in (F–M). ELISA confirmed TGFβ1 secretion at 36 h (N). EMT induction was evaluated by immunoblotting of E-cadherin, vimentin, and N-cadherin (O), with β-actin and Ponceau S as controls and corresponding quantification in (P–R). Data are presented as mean ± SEM (n=3); ***P < 0.001, ****P < 0.0001; NS, not significant (one-way ANOVA).

### BCL2L13 expression shows histology- and site-dependent patterns in human lung cancer

We evaluated BCL2L13 protein expression by IHC in a human lung cancer TMA containing non-cancer lung tissue, primary tumors, and matched lymph-node specimens (Fig. 2A–F). BCL2L13 staining was predominantly cytoplasmic/granular, consistent with its mitochondrial-associated localization. Quantitative TMA analysis showed increased BCL2L13 expression in primary lung tumors relative to non-cancer lung tissue, whereas lymph-node cores showed more heterogeneous staining and lower overall expression across the cohort (Fig. 2B,C).

**Figure 2.**
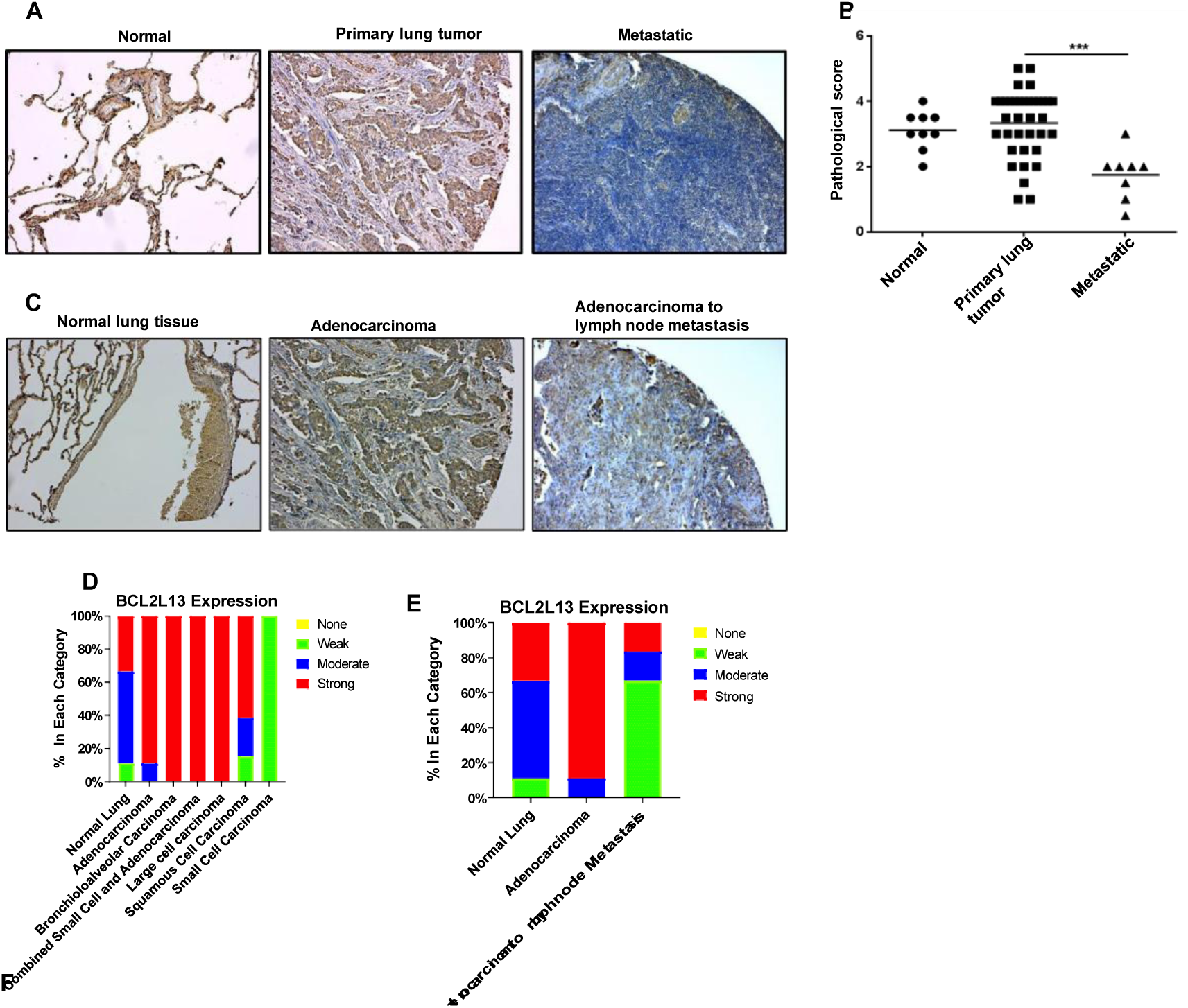
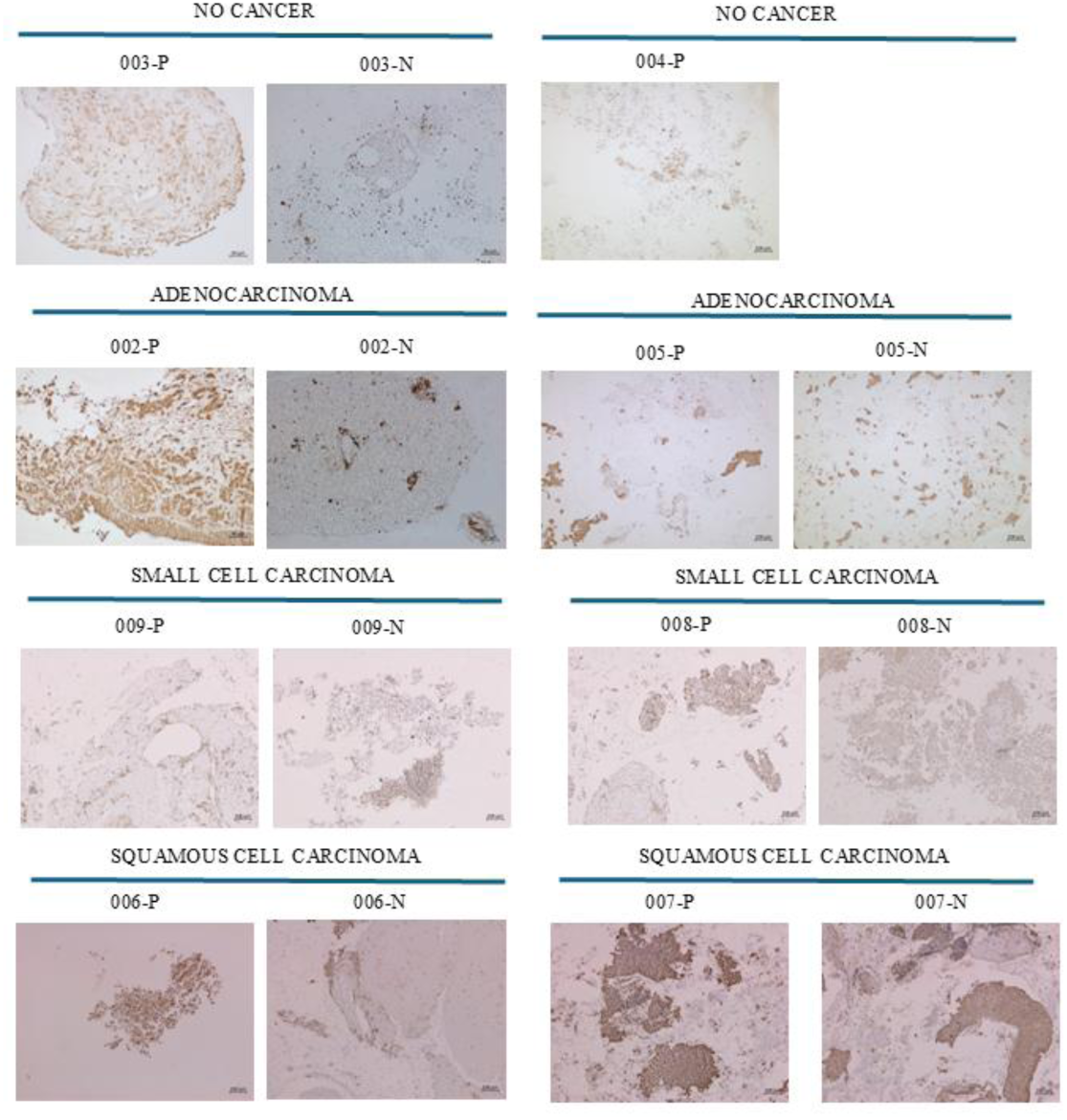
BCL2L13 protein expression varies by lung cancer subtype and tissue compartment in human TMA and matched primary–lymph-node specimens. **(A, C)** Representative BCL2L13 immunohistochemistry (IHC) in human lung cancer tissue microarray sections, including non-cancer lung, primary lung tumors, metastatic tissues, and adenocarcinoma with matched lymph-node metastases. Brown DAB signal indicates BCL2L13 immunoreactivity; nuclei are counterstained with hematoxylin. **(B, E)** Quantification of BCL2L13 staining shows increased expression in primary tumors relative to non-cancer lung tissue, with more heterogeneous and overall lower staining in metastatic cores across the cohort. Subtype analysis indicates lower BCL2L13 expression in small cell carcinoma and higher expression in non-small cell lung cancer subtypes, including large cell carcinoma and adenocarcinoma. **(D)** Distribution of BCL2L13 staining intensity across tissue groups, classified as strong, moderate, weak, or absent. **(F)** Representative paired human patient samples showing BCL2L13 staining in non-cancer lung/lymph-node controls and lung cancer subtypes. P denotes primary lung tissue/tumor and N denotes lymph-node tissue from the same case. Non-cancer lung shows weak-to-patchy basal staining, whereas non-cancer lymph node lacks tumor-associated signal. Adenocarcinoma and squamous cell carcinoma show predominantly cytoplasmic/granular BCL2L13 positivity in primary tumor cells, with retained staining in matched lymph-node metastases when metastatic tumor is morphologically evident. Small cell carcinoma shows weaker and more heterogeneous BCL2L13 staining in tumor-rich regions. Scattered pigment, necrotic or keratinized material, macrophage-associated signal, and nonspecific punctate DAB deposits were excluded from interpretation. Scale bars are shown in individual panels.

Subtype-stratified analysis revealed marked histology-dependent variation. BCL2L13 expression was lowest in small cell carcinoma and highest in NSCLC subtypes, particularly large cell carcinoma, bronchioloalveolar carcinoma, and adenocarcinoma (Fig. 2D). In adenocarcinoma, BCL2L13 expression showed a compartment-specific pattern, with matched lymph-node metastases retaining or increasing tumor-cell staining in selected cases compared with the corresponding primary tumor cores (Fig. 2E).

Representative human patient samples supported the TMA-based interpretation (Fig. 2F). Non-cancer lung showed only weak, patchy basal BCL2L13 staining, while non-cancer lymph-node tissue lacked tumor-associated signal. In adenocarcinoma, primary tumors showed clear cytoplasmic/granular BCL2L13 positivity, and matched nodal metastases retained BCL2L13 expression in malignant glandular deposits. Squamous cell carcinomas showed moderate-to-strong BCL2L13 staining in viable primary tumor nests, with retained expression in nodal metastases when metastatic squamous carcinoma was morphologically evident. In contrast, small cell carcinoma displayed lower and more heterogeneous BCL2L13 staining in both primary and nodal tumor-rich regions.

Clinicopathological correlation analysis across the NSCLC cohort showed no significant association between BCL2L13 expression and stage, tumor type, lymph-node involvement, age, or sex (Table 1). When analyzed by histological subtype relative to non-cancer lung, significant associations were detected in large cell carcinoma and bronchioloalveolar carcinoma (Tables 2 and 3), but not in adenocarcinoma or squamous cell carcinoma (Tables 4 and 5). Together, these data indicate that BCL2L13 expression is enriched in human lung cancer in a subtype-dependent manner, is generally stronger in NSCLC than in small cell carcinoma, and can be retained in confirmed lymph-node metastases. Tumor-specific scoring was restricted to viable carcinoma deposits, with pigment, necrosis, keratinized debris, macrophage-associated signal, and nonspecific punctate DAB deposits excluded from interpretation.

**Table 1.**
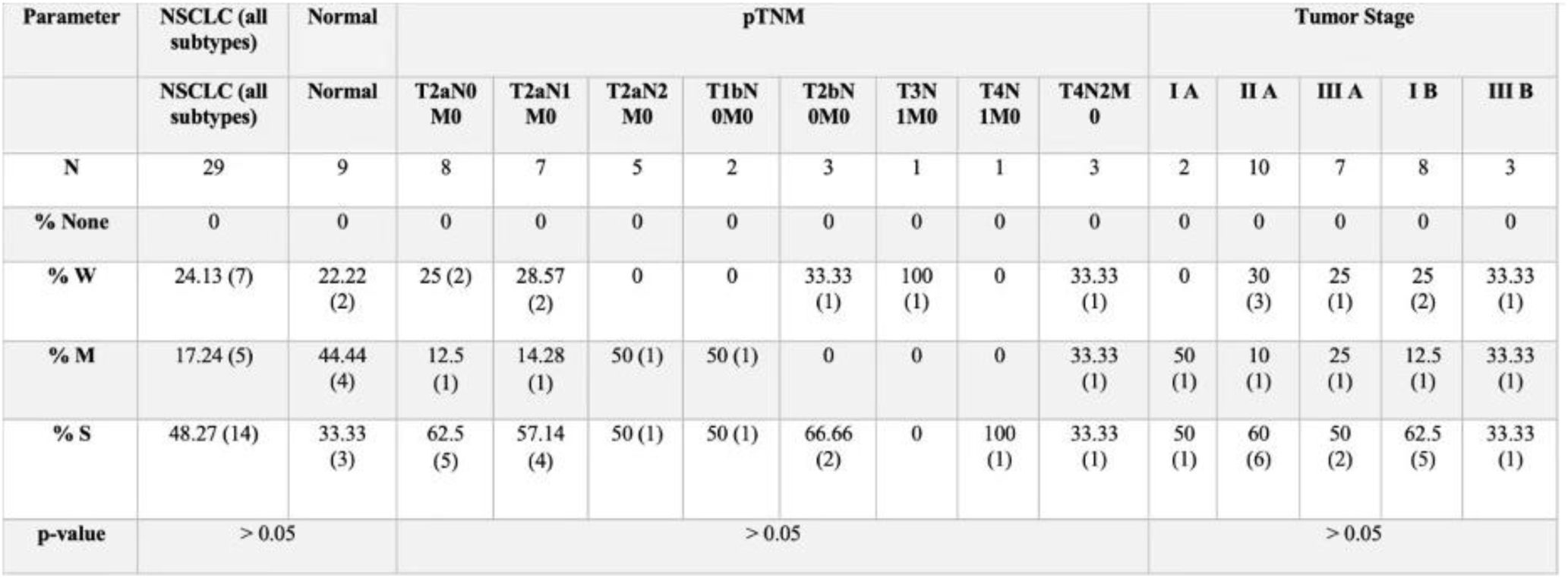
Correlation of BCL2L13 protein expression with clinicopathological features of NSCLC. Pathological Scoring of BCL2L13 expression has been done blindly by three independent cancer pathologists: N: Number of patients in each group, W: Weak expression, M: Medium expression, S: Strong expression.

| Parameter | NSCLC (all subtypes) | Normal | pTNM |  |  |  |  |  |  |  | Tumor Stage |  |  |  |  |
| --- | --- | --- | --- | --- | --- | --- | --- | --- | --- | --- | --- | --- | --- | --- | --- |
|  | NSCLC (all subtypes) | Normal | T2aN0 M0 | T2aN1 M0 | T2aN2 M0 | T1bN 0M0 | T2bN 0M0 | T3N 1M0 | T4N 1M0 | T4N2M 0 | I A | II A | III A | I B | III B |
| N | 29 | 9 | 8 | 7 | 5 | 2 | 3 | 1 | 1 | 3 | 2 | 10 | 7 | 8 | 3 |
| % None | 0 | 0 | 0 | 0 | 0 | 0 | 0 | 0 | 0 | 0 | 0 | 0 | 0 | 0 | 0 |
| % W | 24.13 (7) | 22.22 (2) | 25 (2) | 28.57 (2) | 0 | 0 | 33.33 (1) | 100 (1) | 0 | 33.33 (1) | 0 | 30 (3) | 25 (1) | 25 (2) | 33.33 (1) |
| % M | 17.24 (5) | 44.44 (4) | 12.5 (1) | 14.28 (1) | 50 (1) | 50 (1) | 0 | 0 | 0 | 33.33 (1) | 50 (1) | 10 (1) | 25 (1) | 12.5 (1) | 33.33 (1) |
| % S | 48.27 (14) | 33.33 (3) | 62.5 (5) | 57.14 (4) | 50 (1) | 50 (1) | 66.66 (2) | 0 | 100 (1) | 33.33 (1) | 50 (1) | 60 (6) | 50 (2) | 62.5 (5) | 33.33 (1) |
| p-value | > 0.05 |  | > 0.05 |  |  |  |  |  |  |  | > 0.05 |  |  |  |  |

**Table 2.** Correlation of BCL2L13 protein expression with clinicopathological features of large cell carcinoma. Pathological Scoring of BCL2L13 expression has been done blindly by three independent cancer pathologists: N: Number of patients in each group, W: Weak expression, M: Medium expression, S: Strong expression.

| Parameter | NSCLC subtype | Normal | pTNM |  |  |  |  |  |  |  | Tumor Stage |  |  |  |  |
| --- | --- | --- | --- | --- | --- | --- | --- | --- | --- | --- | --- | --- | --- | --- | --- |
|  | large cell carcinoma | Normal | T2aN0 M0 | T2aN1 M0 | T2aN2 M0 | T1bN0 M0 | T2bN0 M0 | T3N 1M0 | T4N 1M0 | T4N2M 0 | I A | II A | III A | I B | III B |
| N | 3 | 9 | 0 | 1 | 1 | 0 | 1 | 0 | 0 | 0 | 0 | 2 | 1 | 0 | 0 |
| % None | 0 | 0 | 0 | 0 | 0 | 0 | 0 | 0 | 0 | 0 | 0 | 0 | 0 | 0 | 0 |
| % W | 0 | 22.22 (2) | 0 | 0 | 0 | 0 | 0 | 0 | 0 | 0 | 0 | 0 | 0 | 0 | 0 |
| % M | 0 | 44.44 (4) | 0 | 0 | 0 | 0 | 0 | 0 | 0 | 0 | 0 | 0 | 0 | 0 | 0 |
| % S | 100 (3) | 33.33 (3) | 0 | 100 (1) | 100 (1) | 0 | 100 (1) | 0 | 0 | 0 | 0 | 100 (2) | 100 (1) | 0 | 0 |
| p-value | < 0.05 |  | NA |  |  |  |  |  |  |  | NA |  |  |  |  |

**Table 3.** Correlation of BCL2L13 protein expression with clinicopathological features of bronchioloalveolar carcinoma. Pathological Scoring of BCL2L13 expression has been done blindly by three independent cancer pathologists: N: Number of patients in each group, W: Weak expression, M: Medium expression, S: Strong expression.

| Parameter | NSCLC subtype | Normal Muscle | pTNM |  |  |  |  |  |  |  | Tumor Stage |  |  |  |  |
| --- | --- | --- | --- | --- | --- | --- | --- | --- | --- | --- | --- | --- | --- | --- | --- |
|  | bronchioloal veolar carcinoma | Normal | T2aN0 M0 | T2aN1 M0 | T2aN2 M0 | T1bN0 M0 | T2bN0 M0 | T3N 1M0 | T4N 1M0 | T4N2 M0 | I A | II A | III A | I B | III B |
| N | 4 | 9 | 2 | 0 | 0 | 1 | 1 | 0 | 0 | 0 | 1 | 0 | 0 | 3 | 0 |
| % None | 0 | 0 | 0 | 0 | 0 | 0 | 0 | 0 | 0 | 0 | 0 | 0 | 0 | 0 | 0 |
| % W | 0 | 22.22 (2) | 0 | 0 | 0 | 0 | 0 | 0 | 0 | 0 | 0 | 0 | 0 | 0 | 0 |
| % M | 0 | 44.44 (4) | 0 | 0 | 0 | 0 | 0 | 0 | 0 | 0 | 0 | 0 | 0 | 0 | 0 |
| % S | 100 (4) | 33.33 (3) | 100 (2) | 0 | 0 | 100 (1) | 100 (1) | 0 | 0 | 0 | 100 (1) | 0 | 0 | 100 (3) | 0 |
| p-value | < 0.05 |  | NA |  |  |  |  |  |  |  | NA |  |  |  |  |

**Table 4.** Correlation of BCL2L13 protein expression with clinicopathological features of adenocarcinoma. Pathological Scoring of BCL2L13 expression has been done blindly by three independent cancer pathologists: N: Number of patients in each group, W: Weak expression, M: Medium expression, S: Strong expression.

| Parameter | NSCLC subtype | Normal | pTNM |  |  |  |  |  |  |  | Tumor Stage |  |  |  |  |
| --- | --- | --- | --- | --- | --- | --- | --- | --- | --- | --- | --- | --- | --- | --- | --- |
|  | adenocarcinoma | Normal | T2aN0 M0 | T2aN1 M0 | T2aN2 M0 | T1bN 0M0 | T2bN 0M0 | T3N 1M0 | T4N 1M0 | T4N2M 0 | I A | II A | III A | I B | III B |
| N | 7 | 9 | 2 | 2 | 0 | 1 | 1 | 0 | 0 | 1 | 1 | 3 | 1 | 2 | 1 |
| % None | 0 | 0 | 0 | 0 | 0 | 0 | 0 | 0 | 0 | 0 | 0 | 0 | 0 | 0 | 0 |
| % W | 0 | 22.22 (2) | 0 | 0 | 0 | 0 | 0 | 0 | 0 | 0 | 0 | 0 | 100 (1) | 0 | 0 |
| % M | 14.28 (1) | 44.44 (4) | 0 | 0 | 0 | 100 (1) | 0 | 0 | 0 | 0 | 100 (1) | 0 | 0 | 0 | 0 |
| % S | 85.71 (6) | 33.33 (3) | 100 (2) | 100 (2) | 0 | 0 | 100 (1) | 0 | 0 | 100 (1) | 0 | 100 (3) | 0 | 100 (2) | 100 (1) |
| p-value | > 0.05 |  | > 0.05 |  |  |  |  |  |  |  | > 0.05 |  |  |  |  |

**Table 5.** Correlation of BCL2L13 protein expression with clinicopathological features of squamous cell carcinoma. Pathological Scoring of BCL2L13 expression has been done blindly by three independent cancer pathologists: N: Number of patients in each group, W: Weak expression, M: Medium expression, S: Strong expression.

| Parameter | NSCLC subtype | Normal | pTNM |  |  |  |  |  |  |  | Tumor Stage |  |  |  |  |
| --- | --- | --- | --- | --- | --- | --- | --- | --- | --- | --- | --- | --- | --- | --- | --- |
|  | squamous cell carcinoma | Normal | T2aN0 M0 | T2aN1 M0 | T2aN2 M0 | T1bN 0M0 | T2bN 0M0 | T3N 1M0 | T4N 1M0 | T4N2M 0 | I A | II A | III A | I B | III B |
| N | 13 | 9 | 3 | 4 | 1 | 0 | 1 | 1 | 1 | 2 | 0 | 5 | 1 | 3 | 2 |
| % None | 0 | 0 | 0 | 0 | 0 | 0 | 0 | 0 | 0 | 0 | 0 | 0 | 0 | 0 | 0 |
| % W | 53.84 (7) | 22.22 (2) | 66.66 (2) | 50 (2) | 0 | 0 | 100 (1) | 100 (1) | 0 | 50 (1) | 0 | 60 (3) | 100 (1) | 66.33 (2) | 50 (1) |
| % M | 30.76 (4) | 44.44 (4) | 33.33 (1) | 25 (1) | 100 (1) | 0 | 0 | 0 | 0 | 50 (1) | 0 | 20 (1) | 0 | 33.33 (1) | 50 (1) |
| % S | 15.38 (2) | 33.33 (3) | 0 | 25 (1) | 0 | 0 | 0 | 0 | 100 (1) | 0 | 0 | 20 (1) | 0 | 0 | 0 |
| p-value | > 0.05 |  | > 0.05 |  |  |  |  |  |  |  | > 0.05 |  |  |  |  |

### BCL2L13 supports TGFβ1-induced mitophagy and limits EMT-associated plasticity in NSCLC adenocarcinoma cells

Because BCL2L13 increased at mitochondria during TGFβ1-induced mitophagy and EMT (Fig. 1; Suppl. Fig. 1), we next tested whether BCL2L13 functionally contributes to these responses. Stable *BCL2L13* knockdown (KD) and overexpression (OE) models were generated in A549 and LLC cells (Fig. 3A–D; Suppl. Fig. 2A–D), and mitophagy and EMT were assessed using established imaging, fractionation, immunoblotting, and real-time migration assays ^11^. *BCL2L13*-KD reduced TGFβ1-induced mitophagy in both cell lines, as shown by decreased LC3β– mitochondria colocalization and reduced TOMM20–LAMP1 overlap (Fig. 3E–H; Suppl. Fig. 2E–H). Consistently, mitochondrial fractionation demonstrated reduced mitochondrial LC3-II accumulation and attenuated degradation of p62 and TOMM20 after TGFβ1 treatment, whereas BCL2L13-OE enhanced these mitophagy-associated changes (Fig. 3I–P; Suppl. Fig. 2I–P). BCL2L13 manipulation did not substantially alter BNIP3 or NIX localization, including monomeric or dimeric forms, suggesting that the observed mitophagy phenotype was not explained by compensatory changes in these receptors under the conditions tested (61) (Fig. 3I,J,Q–X; Suppl. Fig. 2I,J,Q–X). *BCL2L13* loss was accompanied by a stronger EMT phenotype after TGFβ1 exposure, with increased vimentin and N-cadherin and reduced E-cadherin expression in A549 and LLC cells (Fig. 3Y–BB; Suppl. Fig. 2Y–BB). Conversely, *BCL2L13*-OE partially opposed these EMT-associated changes, reducing mesenchymal-marker expression and preserving epithelial features (Fig. 3Y,DD; Suppl. Fig. 2Y,CC–EE). Functionally, *BCL2L13*-KD increased A549 cell migration in real-time transwell assays (Fig. 3FF,GG). Together, these data support a model in which BCL2L13 facilitates TGFβ1-associated mitophagic processing while restraining EMT-linked phenotypic plasticity in NSCLC adenocarcinoma cells.

**Figure 3.**
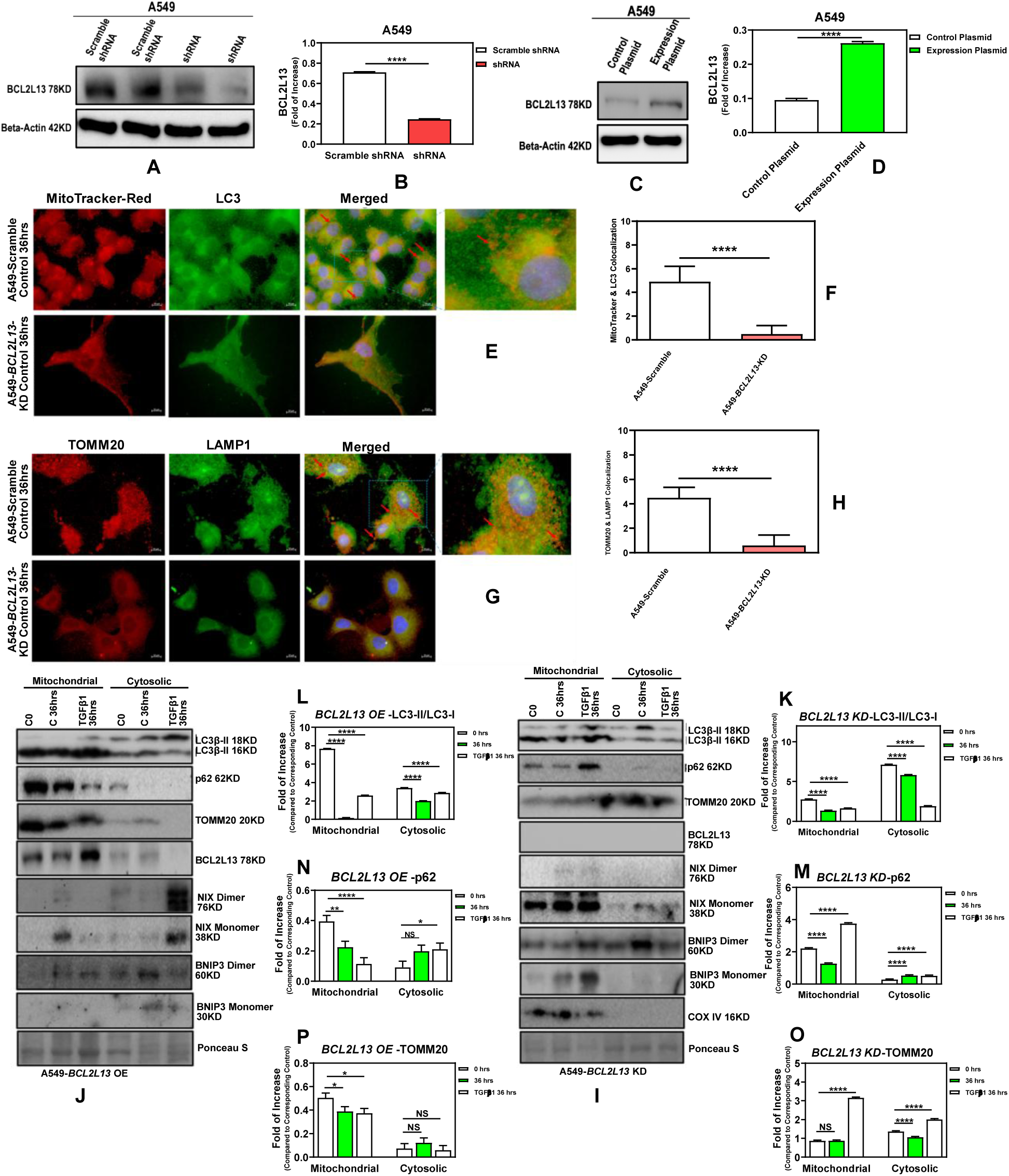
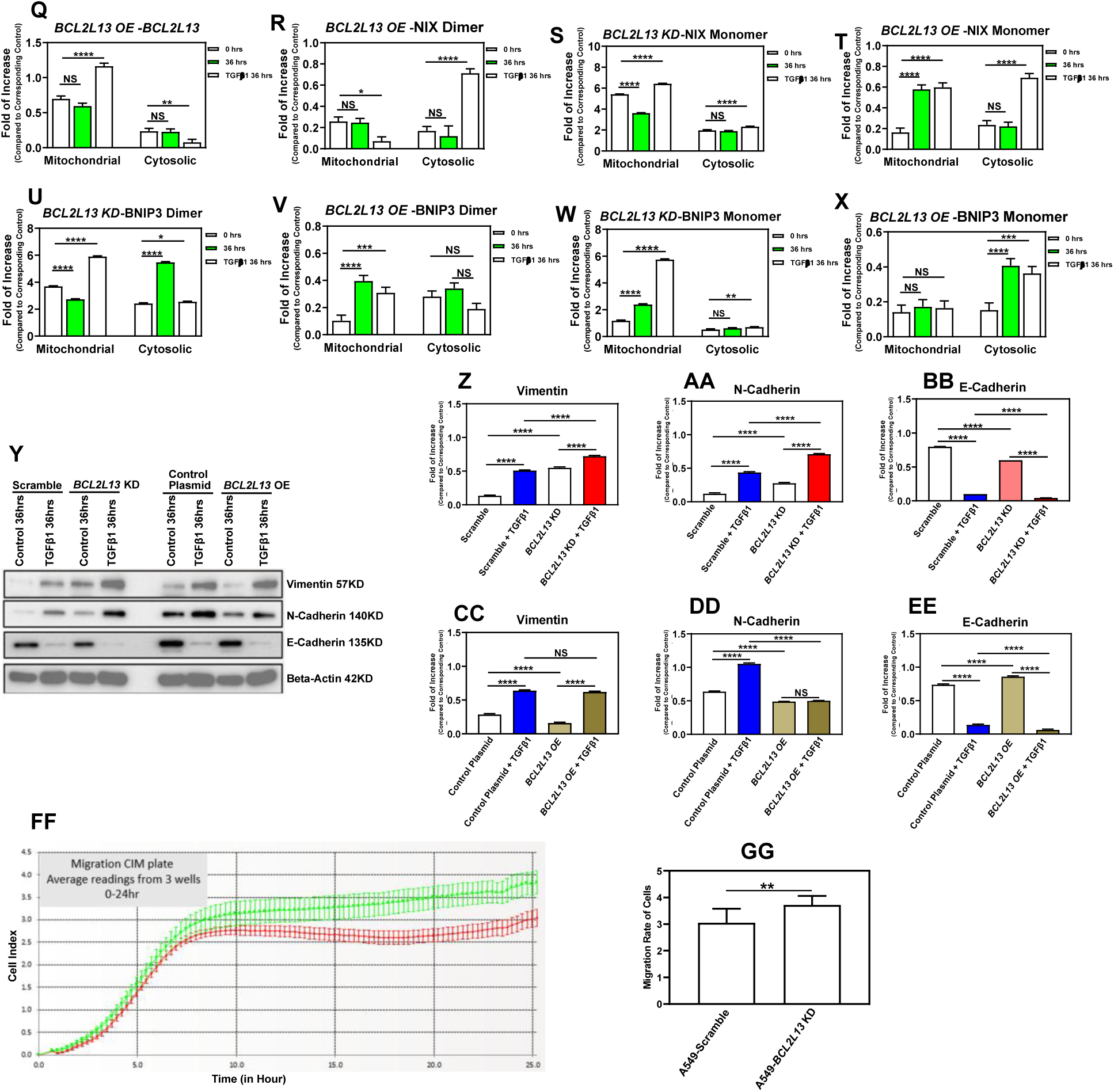
BCL2L13 supports mitophagy-associated responses and limits EMT-linked migration in A549 cells. (A–D) Stable *BCL2L13* knockdown (KD) and overexpression (OE) in A549 cells were validated by immunoblotting. Scramble shRNA and empty pcDNA3 vector were used as controls; β-actin served as loading control. **(E–H)** Following TGFβ1 treatment, *BCL2L13*-KD reduced LC3β–mitochondria colocalization and TOMM20–LAMP1 overlap, consistent with reduced mitophagy-associated trafficking. **(I–P)** Subcellular fractionation showed that *BCL2L13*-KD attenuated mitochondrial LC3-II accumulation and reduced p62 and TOMM20 turnover, whereas *BCL2L13*-OE produced the reciprocal trend after TGFβ1 exposure. COX IV and Ponceau S were used to assess fraction loading/purity. **(Q–X)** BNIP3 and NIX distribution between mitochondrial and cytosolic fractions was not markedly altered by *BCL2L13*-KD or -OE. **(Y–EE)** *BCL2L13*-KD enhanced TGFβ1-associated EMT marker changes, including increased vimentin and N-cadherin and reduced E-cadherin, whereas *BCL2L13*-OE partially opposed these changes. **(FF, GG)** Real-time migration analysis showed increased migration in *BCL2L13*-KD cells. Cells were treated with TGFβ1, 5 ng/ml, for 36 h where indicated. Scale bars, 10 μm. Data are mean ± SEM from three independent experiments, unless otherwise indicated; imaging quantification was performed from 10 fields with approximately five cells per field. NS, not significant; *p ≤ 0.05, **p ≤ 0.01, ***p ≤ 0.001, ****p ≤ 0.0001.

### BCL2L13 modulates mitochondrial bioenergetics and functional parameters in NSCLC adenocarcinoma cells

Building on the finding that BCL2L13 contributes to mitophagy-associated responses in NSCLC adenocarcinoma cells (Fig. 3; Suppl. Fig. 2), we next tested whether BCL2L13 also affects mitochondrial function, consistent with the established involvement of BCL2 family proteins in mitochondrial regulation ^28–30^. Seahorse analysis showed that *BCL2L13* knockdown reduced mitochondrial oxygen consumption in A549 cells, with decreases in basal respiration, maximal respiration, spare respiratory capacity, ATP-linked respiration, and non-mitochondrial respiration compared with control cells (Fig. 4A–F). *BCL2L13* overexpression also reduced several respiratory parameters relative to vector control, indicating that altered *BCL2L13* expression perturbs mitochondrial respiration in a dosage- and context-dependent manner rather than producing a simple gain- or loss-of-function effect.

**Figure 4.**
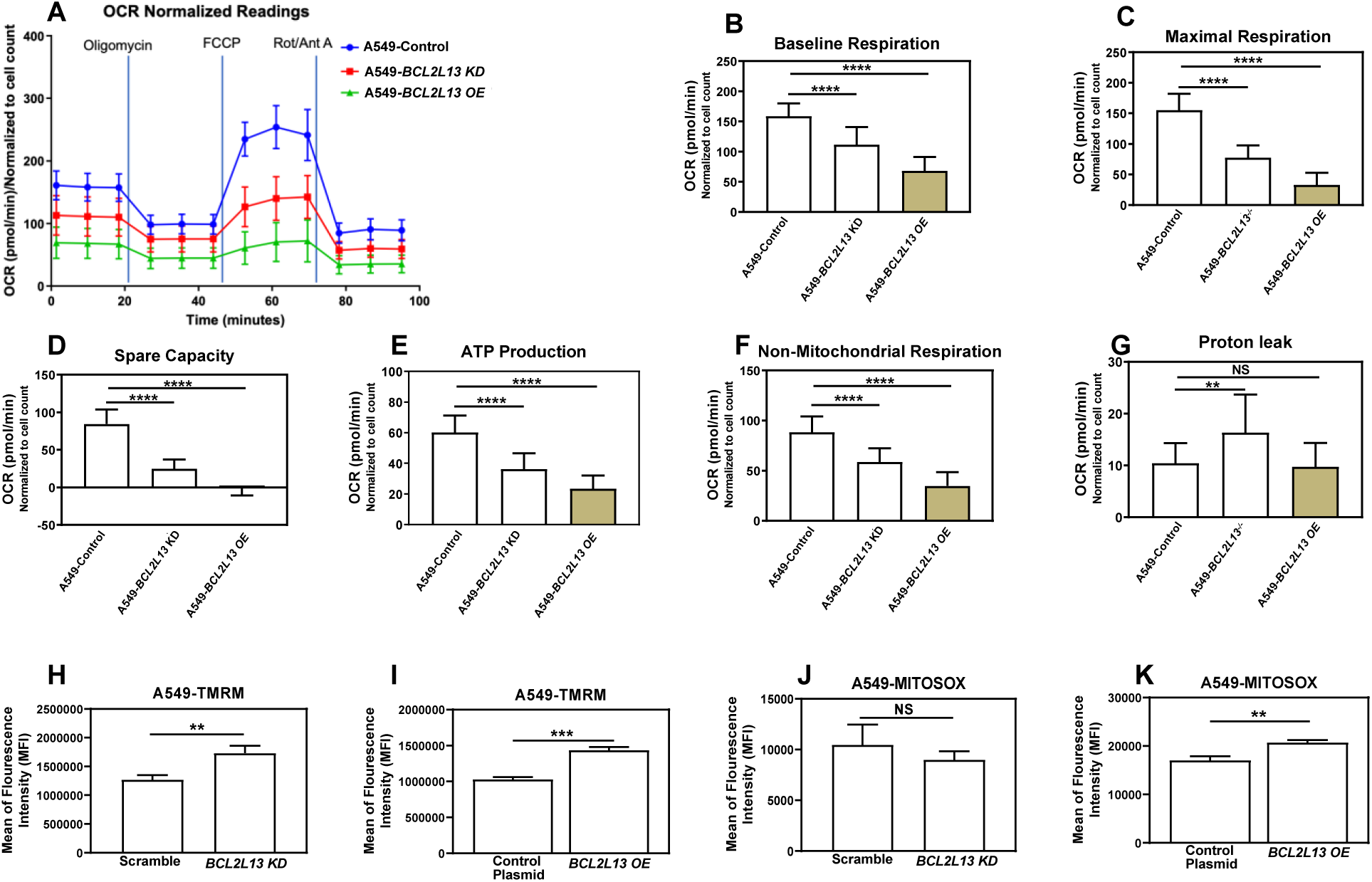
BCL2L13 alters mitochondrial respiratory capacity, membrane potential and mitochondrial ROS in A549 cells. **(A)** Seahorse XF oxygen-consumption rate (OCR) profile in A549 control, *BCL2L13* knockdown (KD) and *BCL2L13* overexpression (OE) cells following sequential addition of oligomycin, FCCP, rotenone and antimycin A. **(B–F)** Quantification of respiratory parameters showed reduced basal respiration, maximal respiration, spare respiratory capacity, ATP-linked respiration and non-mitochondrial respiration after *BCL2L13*-KD, with similar suppression of several OCR-derived parameters after *BCL2L13*-OE. **(G)** Proton leak was increased in *BCL2L13*-KD cells, whereas *BCL2L13*-OE cells did not show a significant change relative to vector control. **(H,I)** TMRM fluorescence showed increased mitochondrial membrane potential signal in both *BCL2L13*-KD and *BCL2L13*-OE cells. **(J,K)** MitoSOX analysis showed no significant change in mitochondrial ROS after *BCL2L13*-KD, but increased mitochondrial ROS after *BCL2L13*-OE. OCR values were normalized to cell count. Data are mean ± SEM from three independent experiments. NS, not significant; *p ≤ 0.05, **p ≤ 0.01, ***p ≤ 0.001, ****p ≤ 0.0001.

*BCL2L13* knockdown increased proton leak, whereas this effect was not clearly observed with *BCL2L13* overexpression (Fig. 4G). These data suggest that loss of *BCL2L13* compromises respiratory efficiency and reduces the capacity of A549 cells to increase mitochondrial respiration under stress. However, because both KD and OE decreased OCR-related parameters, the data are best interpreted as evidence that balanced *BCL2L13* expression is required for mitochondrial bioenergetic homeostasis.

We next assessed mitochondrial membrane potential and mitochondrial ROS. TMRM staining showed increased fluorescence in both *BCL2L13*-KD and *BCL2L13*-OE cells compared with their respective controls (Fig. 4H,I), suggesting altered mitochondrial membrane potential regulation. MitoSOX analysis showed no significant change in mitochondrial ROS after *BCL2L13* knockdown, whereas *BCL2L13* overexpression increased mitochondrial ROS (Fig. 4J,K). These findings indicate that BCL2L13 influences mitochondrial respiratory capacity, membrane potential, and redox state in A549 NSCLC adenocarcinoma cells ^31–34^.

### BCL2L13 is required for depolarization-induced mitophagy and shapes EMT-associated responses in NSCLC cells

To further test whether BCL2L13 contributes to mitophagy independently of TGFβ1 stimulation, we used CCCP, a mitochondrial depolarizing agent commonly used to induce mitophagy ^35^. MTT-based dose optimization identified CCCP conditions that preserved sufficient viability while inducing mitophagy-associated responses in scramble and *BCL2L13*-KD A549 and LLC cells (Fig. 5A,B; Suppl. Fig. 3A,B).

**Figure 5.**
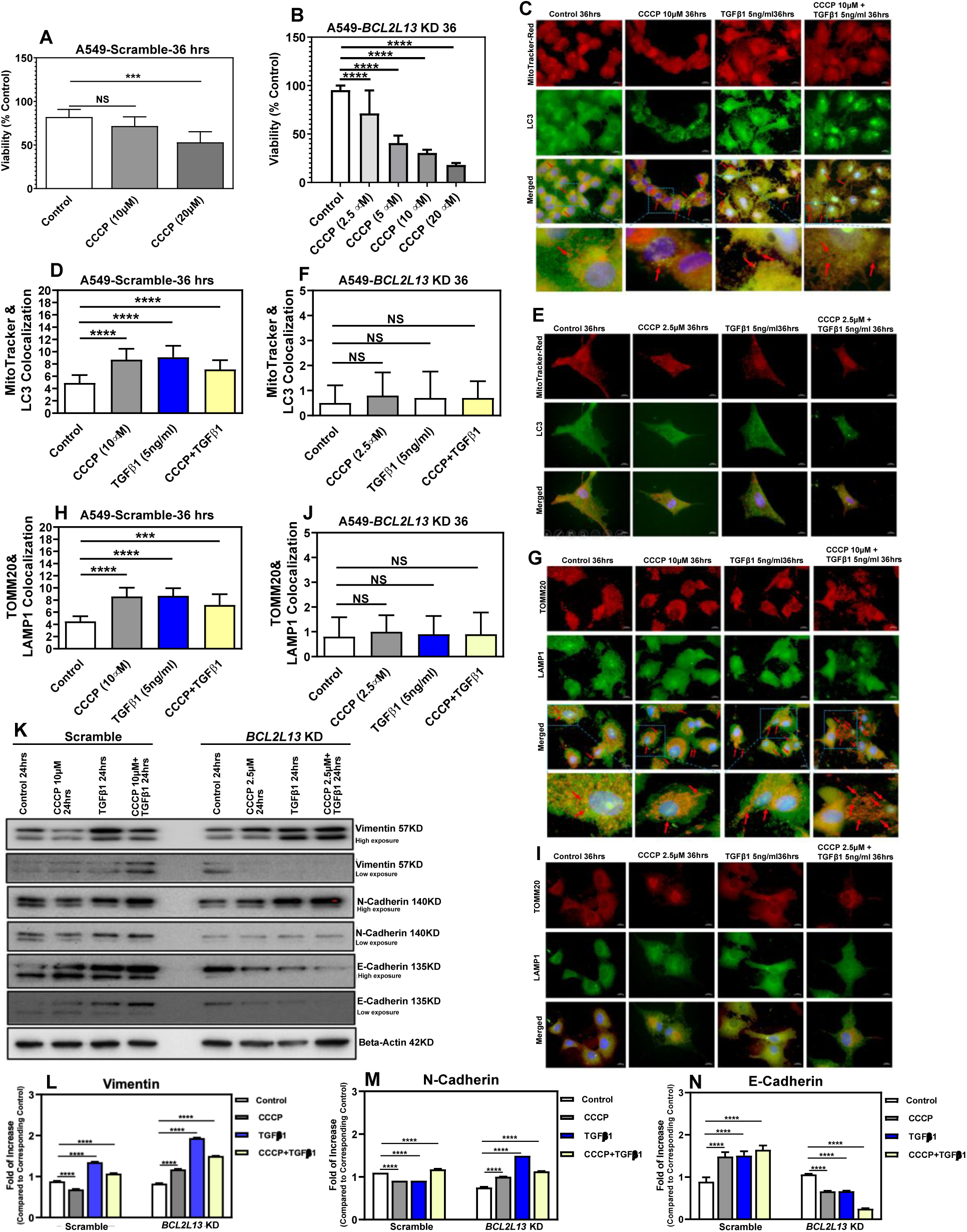
BCL2L13 depletion attenuates CCCP-associated mitophagy and alters EMT marker responses in A549 cells. (A,B) MTT-based dose optimization of CCCP in scramble and *BCL2L13*-KD A549 cells after 48 h treatment. CCCP concentrations of 10 µM for scramble cells and 2.5 µM for *BCL2L13*-KD cells were selected for subsequent experiments based on preserved viability. **(C–J)** Immunocytochemical analysis after CCCP, TGFβ1, or combined CCCP/TGFβ1 treatment showed increased LC3β–mitochondria colocalization and TOMM20–LAMP1 overlap in scramble cells, whereas these mitophagy-associated signals were reduced in *BCL2L13*-KD cells. **(K–N)** Immunoblot analysis showed that CCCP decreased vimentin and N-cadherin and increased E-cadherin in scramble cells, while this EMT marker pattern was not reproduced in *BCL2L13*-KD cells. TGFβ1 increased EMT-associated marker changes, with stronger mesenchymal-marker expression in *BCL2L13*-KD cells. β-actin served as loading control. Cells were treated with TGFβ1, 5 ng/ml, where indicated. ICC analyses were performed after 36 h treatment; immunoblot analyses were performed after 24 h treatment. MTT data are mean ± SD from 15 measurements across three independent experiments; remaining data are mean ± SEM from three independent experiments. NS, not significant; *p ≤ 0.05, **p ≤ 0.01, ***p ≤ 0.001, ****p ≤ 0.0001.

In scramble cells, CCCP, TGFβ1, and combined CCCP/TGFβ1 treatment increased LC3β– mitochondria colocalization and TOMM20–LAMP1 overlap in both A549 and LLC cells, consistent with induction of mitophagy-associated trafficking (Fig. 5C,D,G,H; Suppl. Fig. 3C,D,G,H). These responses were markedly reduced in *BCL2L13*-KD cells (Fig. 5E,F,I,J; Suppl. Fig. 3E,F,I,J), indicating that BCL2L13 is required for efficient CCCP-induced mitophagy under these conditions.

We next asked whether CCCP-associated mitophagy was linked to EMT marker changes. In scramble A549 cells, CCCP treatment reduced vimentin and N-cadherin and increased E-cadherin, suggesting attenuation of EMT-associated features (Fig. 5L–N). This response was not reproduced in *BCL2L13*-KD cells, in which mesenchymal marker expression was maintained or increased. Consistent with our earlier findings, TGFβ1 promoted a stronger EMT marker profile in *BCL2L13*-deficient cells than in scramble controls (Fig. 5L–N; see also Fig. 3). In conclusion, these data indicate that BCL2L13 supports depolarization-induced mitophagy and that loss of *BCL2L13* uncouples CCCP treatment from the attenuation of EMT-associated marker expression. These findings further support a functional link between mitochondrial quality control and epithelial– mesenchymal plasticity in NSC

### BCL2L13 perturbation is associated with sphingolipid remodeling and PI3K–AKT-linked lipid-network signatures

Ceramide metabolism has been linked to mitophagy through ceramide-dependent recruitment of LC3B-II-positive autophagic membranes to mitochondria ^36,37^. Because BCL2L13 has also been reported to interact with CerS2 and CerS6 ^10^, we examined whether altered BCL2L13 expression was associated with changes in sphingolipid composition in A549 NSCLC cells. Lipidomic profiling showed a broad shift in sphingolipid-related species after *BCL2L13* knockdown (KD) or overexpression (OE), with several ceramide-associated classes, including Cer, dhCer, MHC, DHC and THC, separating across experimental groups (Fig. 6A).

**Figure 6.**
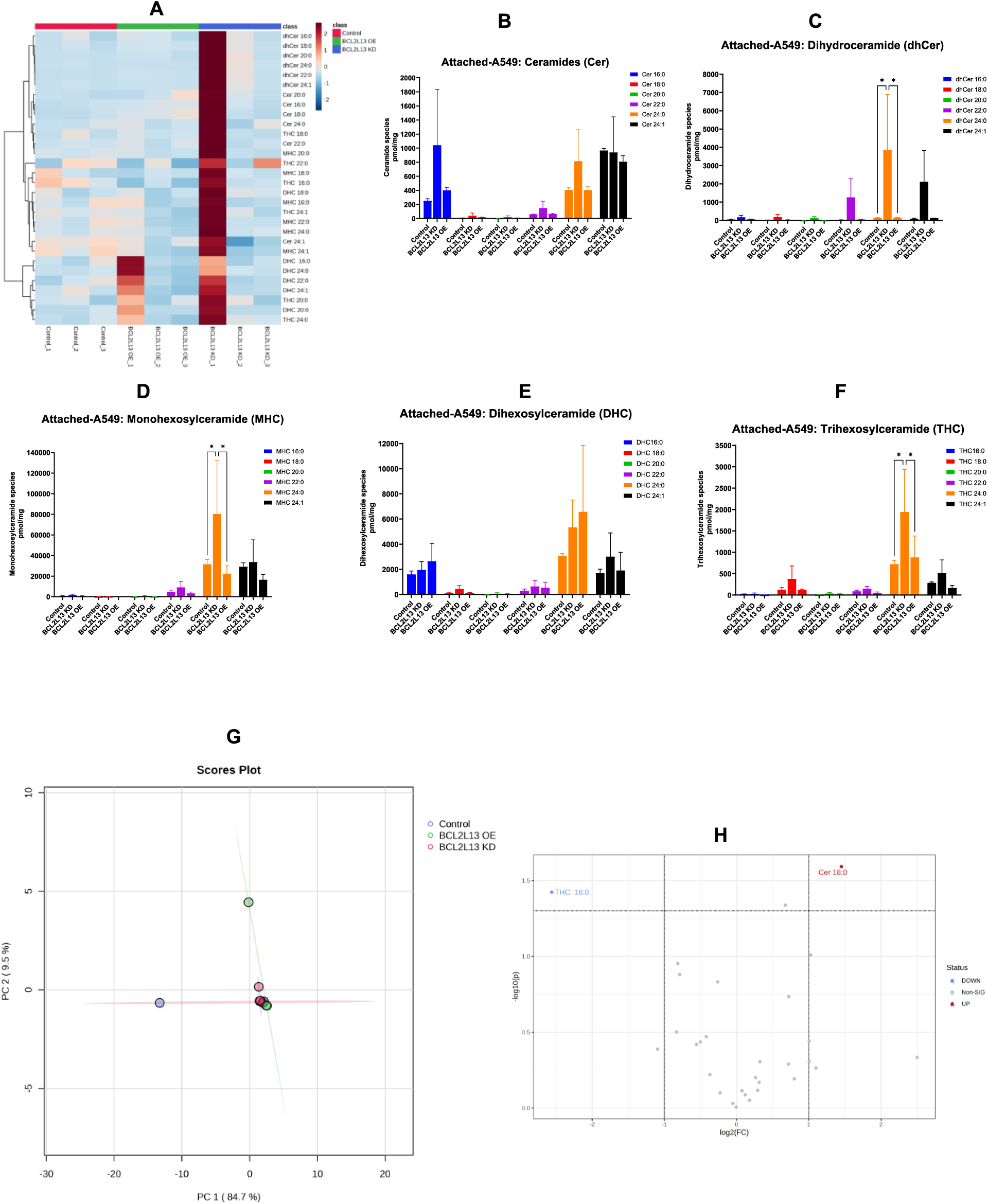
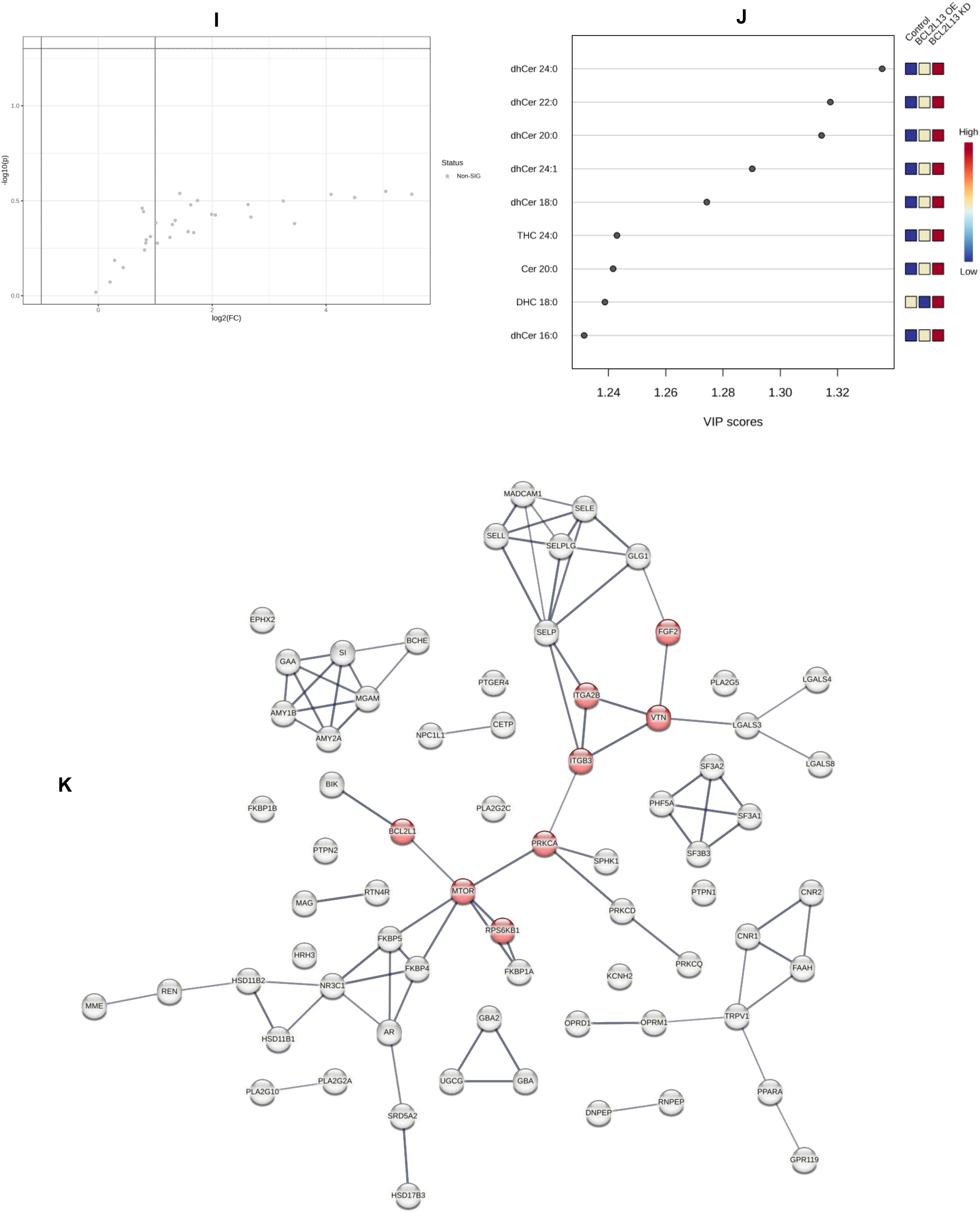
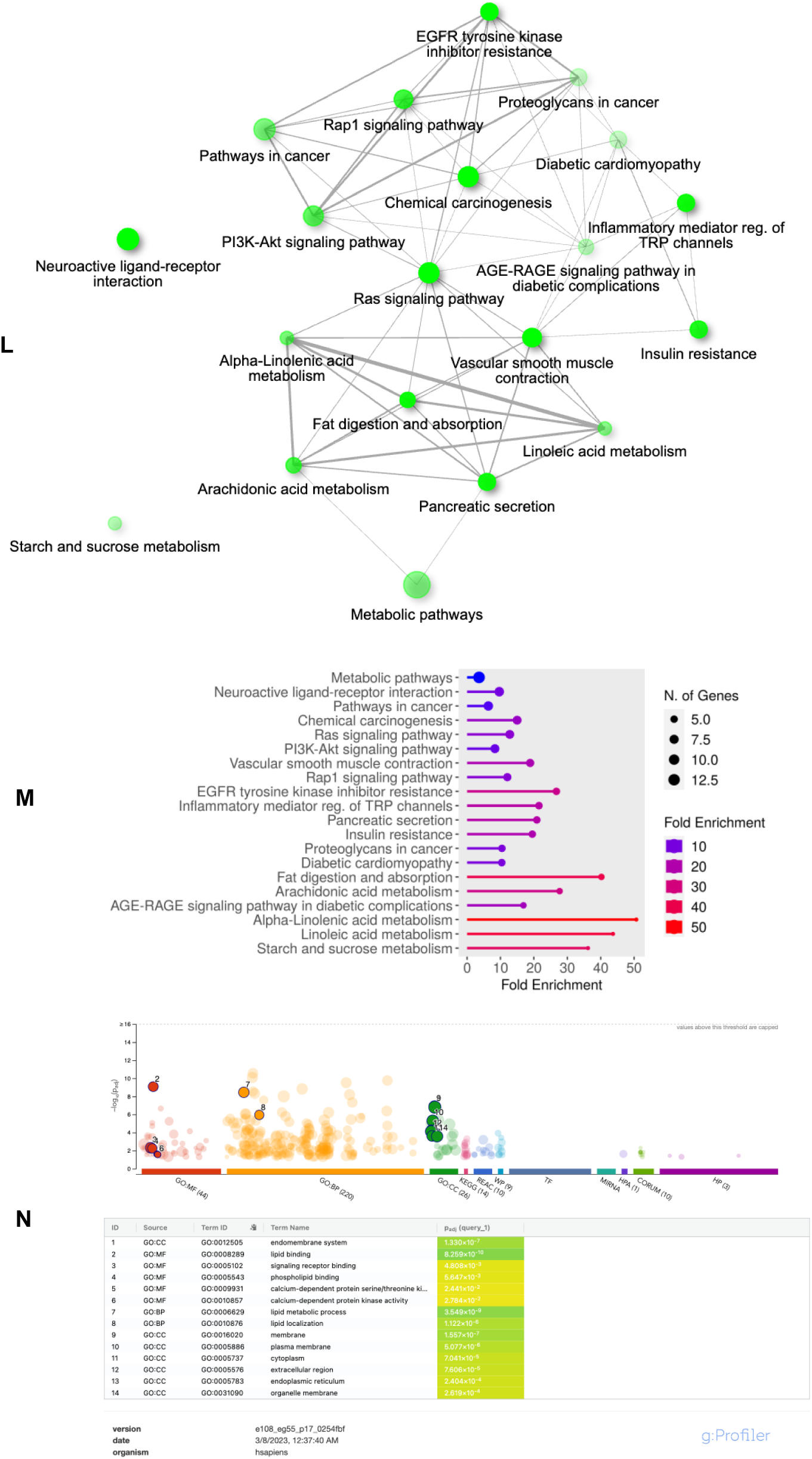
BCL2L13 perturbation is associated with selective sphingolipid remodeling and PI3K–AKT-linked lipid-network signatures in A549 cells. **(A)** Heatmap showing relative abundance of ceramide-related lipid classes, including Cer, dhCer, MHC, DHC and THC, in control, *BCL2L13* knockdown (KD) and *BCL2L13* overexpression (OE) A549 cells. Scramble shRNA and empty-vector cells served as the respective controls. **(B–F)** Quantification of ceramide-family lipid species showing class- and species-selective changes after *BCL2L13*-KD or -OE. **(G)** Principal component analysis of lipidomic profiles from control, KD and OE cells. **(H,I)** Volcano plots showing differentially abundant lipid species in *BCL2L13*-OE versus vector control and *BCL2L13*-KD versus scramble control. **(J)** PLS-DA/VIP analysis identifying lipid species contributing to group separation. **(K)** Predicted protein-interaction network generated from targets associated with prioritized VIP lipids, highlighting PI3K–AKT-related nodes. **(L,M)** KEGG pathway enrichment analysis of lipid-associated target networks. **(N)** Functional enrichment of molecular function, biological process and cellular component terms associated with the prioritized lipid-linked network. Data are mean ± SEM from three independent experiments. NS, not significant; *p ≤ 0.05 by two-way ANOVA.

Class-level analysis did not show uniform changes in total Cer or DHC abundance (Fig. 6B,E). Instead, *BCL2L13*-KD was associated with selective increases in C24-containing species, including dhCer 24:0, MHC 24:0 and THC 24:0 (Fig. 6C,D,F). PCA further separated control, KD and OE samples, indicating that BCL2L13 expression status is associated with a distinct lipidomic profile (Fig. 6G). Differential analysis showed a more restricted pattern, with *BCL2L13*-OE increasing Cer 18:0 and decreasing THC 16:0, whereas KD did not yield statistically significant species-level changes under the same thresholding criteria (Fig. 6H,I).

To prioritize lipid species contributing to group separation, PLS-DA/VIP analysis identified eight candidate lipids with VIP scores >1.2 (Fig. 6J). As an exploratory annotation step, these lipids were analyzed using SwissTargetPrediction and UniProt-based network mapping. The resulting interaction network was enriched for PI3K–AKT-related pathways (Fig. 6K–M), while ShinyGo analysis linked the predicted networks to organelle-associated molecular functions, biological processes and cellular components (Fig. 6N). These analyses suggest that BCL2L13-associated lipid remodeling may intersect with signaling pathways relevant to mitochondrial regulation and oncogenic adaptation.

Concurrently, these data indicate that BCL2L13 perturbation is associated with selective remodeling of sphingolipid-related lipid species rather than a uniform increase or decrease in total ceramide classes. The increase in C24-containing species after *BCL2L13*-KD is consistent with a possible link between BCL2L13 and CerS2/6-associated lipid metabolism ^10^, but the present data do not directly measure CerS2/6 activity or enzyme complex formation. Therefore, the lipidomic and network findings support a model in which BCL2L13 contributes to sphingolipid-state regulation in NSCLC cells, with potential connections to mitophagy and PI3K–AKT-associated signaling that require further biochemical validation.

### BCL2L13 loss attenuates detachment-induced apoptosis in NSCLC adenocarcinoma cells

Resistance to anoikis is an important feature of metastatic dissemination in NSCLC and other cancers ^38,39^. Given previous reports linking BCL2L13 to apoptotic regulation in several cellular contexts ^40–42^, we examined whether BCL2L13 influences detachment-induced apoptosis in NSCLC adenocarcinoma models. Under anchorage-independent conditions, *BCL2L13* knockdown reduced apoptosis in both A549 and LLC cells compared with their corresponding controls (Fig. 7A; Suppl. Fig. 4A). These data indicate that loss of *BCL2L13* increases survival during detachment, consistent with a role for BCL2L13 in limiting anoikis resistance.

**Figure 7.**
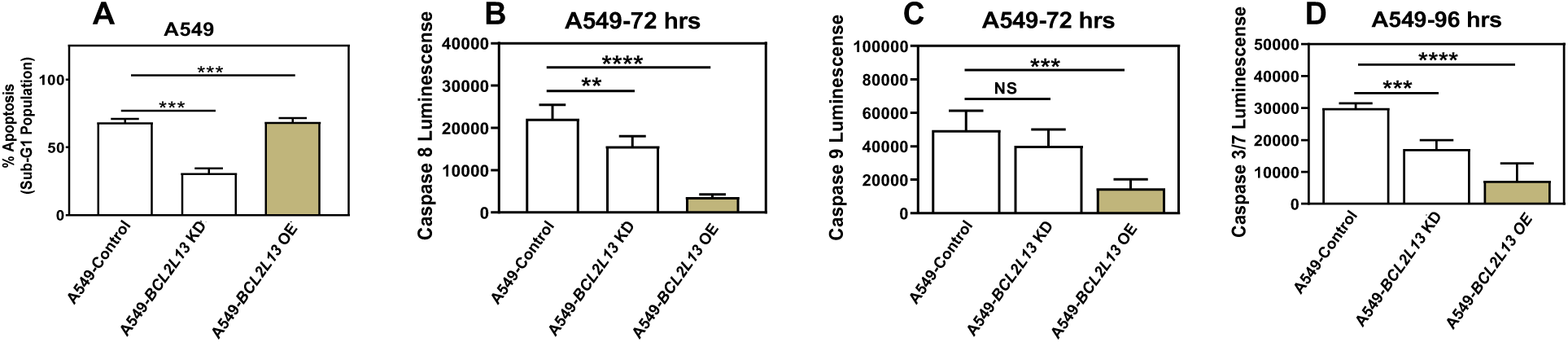
BCL2L13 depletion reduces anoikis-associated apoptosis in A549 cells with non-parallel caspase responses. **(A)** Anoikis-associated apoptosis was assessed in *BCL2L13* knockdown (KD), *BCL2L13* overexpression (OE), and corresponding control A549 cells after 96 h culture under non-adherent conditions using the PI Nicoletti assay. *BCL2L13*-KD reduced detachment-induced apoptosis relative to scramble control, whereas *BCL2L13*-OE increased apoptosis relative to empty-vector control. **(B–D)** Caspase-Glo analysis under anoikis conditions showed reduced caspase-8 and caspase-3/7 activity in *BCL2L13*-KD cells, with a non-significant decrease in caspase-9 activity. *BCL2L13*-OE reduced caspase-8, caspase-9 and caspase-3/7 activity relative to vector control, indicating that caspase activity did not scale directly with the apoptosis phenotype measured by PI staining. Caspase-8 and caspase-9 were measured at 72 h, and caspase-3/7 at 96 h. Data are mean ± SD for the PI assay and mean ± SEM for caspase assays from three independent experiments. NS, not significant; *p ≤ 0.05, **p ≤ 0.01, ***p ≤ 0.001, ****p ≤ 0.0001.

We next measured initiator and executioner caspase activity to determine whether altered apoptosis was accompanied by changes in canonical caspase signaling (Zohny, Zamzami et al. 2019; Başoğlu-Ünal, Becer et al. 2023; Tsai, Chen et al. 2023). In A549 cells, both BCL2L13 knockdown and overexpression reduced caspase-8, caspase-9 and caspase-3/7 activity (Fig. 7B– D). In LLC cells, *BCL2L13* knockdown did not significantly affect caspase activity, whereas BCL2L13 overexpression increased caspase activation (Suppl. Fig. 4B–D). Thus, *BCL2L13* loss consistently reduced detachment-induced apoptosis, but its effect on caspase activity was cell-line and expression-state dependent. Together, these findings suggest that BCL2L13 contributes to anoikis sensitivity in NSCLC adenocarcinoma cells. The reduced apoptosis observed after *BCL2L13* knockdown is not fully explained by uniform changes in initiator or executioner caspase activity, indicating that additional survival or mitochondrial regulatory mechanisms may contribute to detachment tolerance in *BCL2L13*-deficient cells.

### BCL2L13 depletion is associated with altered BCL2-family balance and site-specific FAK phosphorylation during anoikis

Because *BCL2L13* loss reduced detachment-induced apoptosis, we next asked whether this phenotype was accompanied by changes in BCL2-family proteins or focal adhesion kinase (FAK) signaling, both of which regulate anoikis through mitochondrial and adhesion-linked survival pathways ^43,44^ ^45–47^. Detached A549 cells were analyzed after 72 h by immunoblotting for selected pro- and anti-apoptotic BCL2-family members and for total and phosphorylated FAK.

*BCL2L13* knockdown did not produce a uniform reduction in pro-apoptotic proteins. BAX remained largely unchanged, whereas BAK, BNIP3, NIX and the tBID/total BID ratio were increased (Fig. 8A–J). Because tBID reflects caspase-8-dependent BID cleavage ^48^, this increase suggests that upstream apoptotic signaling is not completely suppressed in *BCL2L13*-deficient cells. In parallel, anti-apoptotic proteins were differentially regulated: MCL1 and BCL2 were increased, whereas BCL-XL was reduced (Fig. 8P–S). Thus, the reduced anoikis observed after *BCL2L13* depletion is not explained by a simple global loss of pro-apoptotic signaling, but rather by a mixed remodeling of apoptotic regulators.

**Figure 8.**
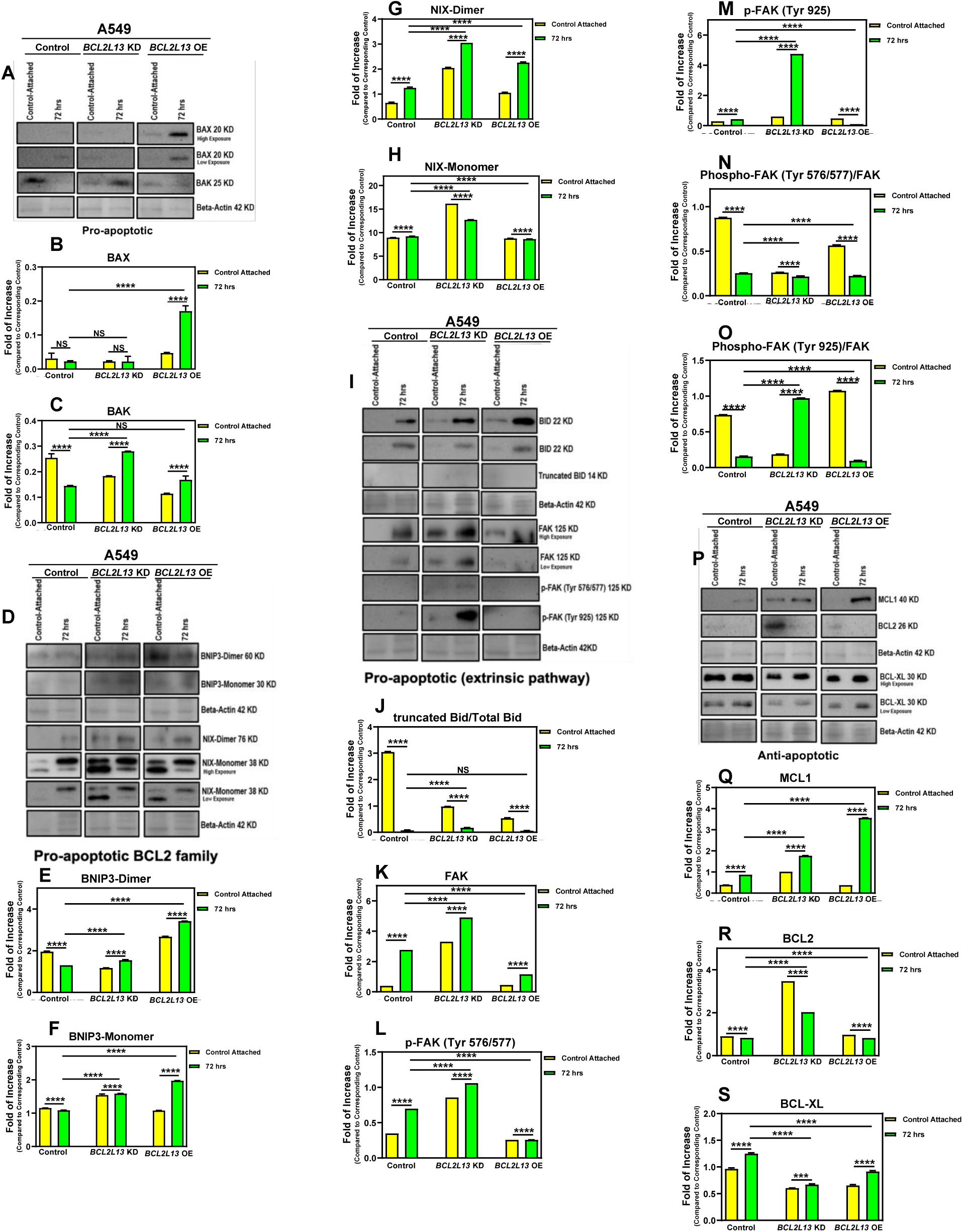
BCL2L13 perturbation alters BCL2-family proteins and FAK phosphorylation during anoikis in A549 cells. *BCL2L13* knockdown (KD), overexpression (OE), and corresponding control A549 cells were cultured under non-adherent conditions for 72 h and analyzed by immunoblotting. Scramble shRNA and empty-vector cells served as KD and OE controls, respectively; attached control cells were included as baseline reference. **(A–H)** Analysis of pro-apoptotic BCL2-family proteins showed selective changes rather than a uniform apoptotic shift: *BCL2L13*-OE increased BAX, *BCL2L13*-KD increased BAK, BNIP3 was increased in both KD and OE conditions, and NIX showed differential regulation between KD and OE cells. **(I–O)** Analysis of BID and FAK signaling showed altered tBID/total BID and site-specific changes in FAK phosphorylation. Total FAK and phospho-FAK levels were modified by *BCL2L13* perturbation, with reduced p-FAK Tyr576/577/FAK ratio and divergent regulation of p-FAK Tyr925/FAK, most notably increased Tyr925 phosphorylation ratio in KD cells and reduced ratio in OE cells. **(P–S)** Anti-apoptotic BCL2-family proteins were also differentially regulated, with increased MCL1 and BCL2 in KD cells and reduced BCL-XL. β-actin served as loading control. Densitometry was performed using AlphaEase FC and GraphPad Prism. Data are mean ± SEM from three independent experiments. NS, not significant; *p ≤ 0.05, **p ≤ 0.01, ***p ≤ 0.001, ****p ≤ 0.0001.

FAK signaling also showed site-specific changes. *BCL2L13*-KD cells displayed reduced phosphorylation of FAK at Tyr576/577, a site associated with FAK catalytic activation downstream of Src-dependent signaling ^49^, while phosphorylation at Tyr925 was increased (Fig. 8I,N,O). This pattern suggests that *BCL2L13* loss alters FAK signaling quality rather than uniformly suppressing or activating the pathway. Reduced Tyr576/577 phosphorylation may reflect diminished canonical adhesion-linked FAK activation, whereas increased Tyr925 phosphorylation may indicate engagement of alternative FAK–Src/MAPK-associated signaling routes (Song, Ye et al. 2018) ^41,50^. Such signaling changes may contribute to detachment tolerance, although they do not by themselves establish causality ^51^. These data show that *BCL2L13* depletion is associated with reduced anoikis, selective remodeling of BCL2-family proteins, and altered FAK phosphorylation under detachment conditions. Because several pro-apoptotic markers were increased despite reduced apoptosis, *BCL2L13*-dependent anoikis sensitivity likely reflects coordinated changes in mitochondrial and adhesion-associated signaling rather than simple suppression of the apoptotic machinery. Given the importance of mitochondrial localization in BCL2-family function during anoikis ^52^, we next examined whether BCL2L13 affects the subcellular distribution of these proteins.

### BCL2L13 depletion alters mitochondrial BCL2-family distribution without reducing pro-apoptotic mitochondrial recruitment during anoikis

Mitochondrial redistribution of BCL2-family proteins is a central step in apoptotic execution ^53,54^. To determine whether the reduced anoikis observed after *BCL2L13* knockdown was associated with altered localization of apoptotic regulators, we performed subcellular fractionation and immunoblotting in A549 cells cultured under detachment conditions for 36 h. BAX was enriched in the mitochondrial fraction after detachment in both scramble and *BCL2L13*-KD cells, whereas BAK showed increased mitochondrial localization specifically in *BCL2L13*-KD cells (Fig. 9A– F). BNIP3 and NIX, including monomeric and dimeric forms, also showed maintained or increased mitochondrial localization in *BCL2L13*-KD cells (Fig. 9G–Q). BCL2L13 itself was enriched in the mitochondrial fraction in control cells, consistent with its mitochondrial association under anoikis conditions (Fig. 9G,L). BID processing showed a different pattern. In scramble cells, detachment increased the tBID/total BID signal in total lysate and cytosolic fractions, whereas mitochondrial tBID was not increased and remained comparatively low (Fig. 9R,S). In *BCL2L13*-KD cells, the tBID/total BID signal was markedly reduced after detachment, particularly in total and cytosolic fractions, with no clear mitochondrial enrichment (Fig. 9T,U). Anti-apoptotic BCL2-family proteins, including MCL1, BCL2 and BCL-XL, also showed protein- and compartment-specific redistribution rather than a uniform shift (Fig. 9V–CC). Together, these data indicate that *BCL2L13* depletion does not reduce mitochondrial recruitment of several canonical pro-apoptotic BCL2-family proteins during anoikis. Instead, the reduced apoptosis observed in *BCL2L13*-KD cells is associated with altered BID cleavage/redistribution and broader changes in compartment-specific BCL2-family signaling. This suggests that anoikis resistance in *BCL2L13*-deficient cells is unlikely to result simply from impaired mitochondrial apoptotic priming, but may involve altered apoptotic execution downstream of mitochondrial recruitment.

**Figure 9.**
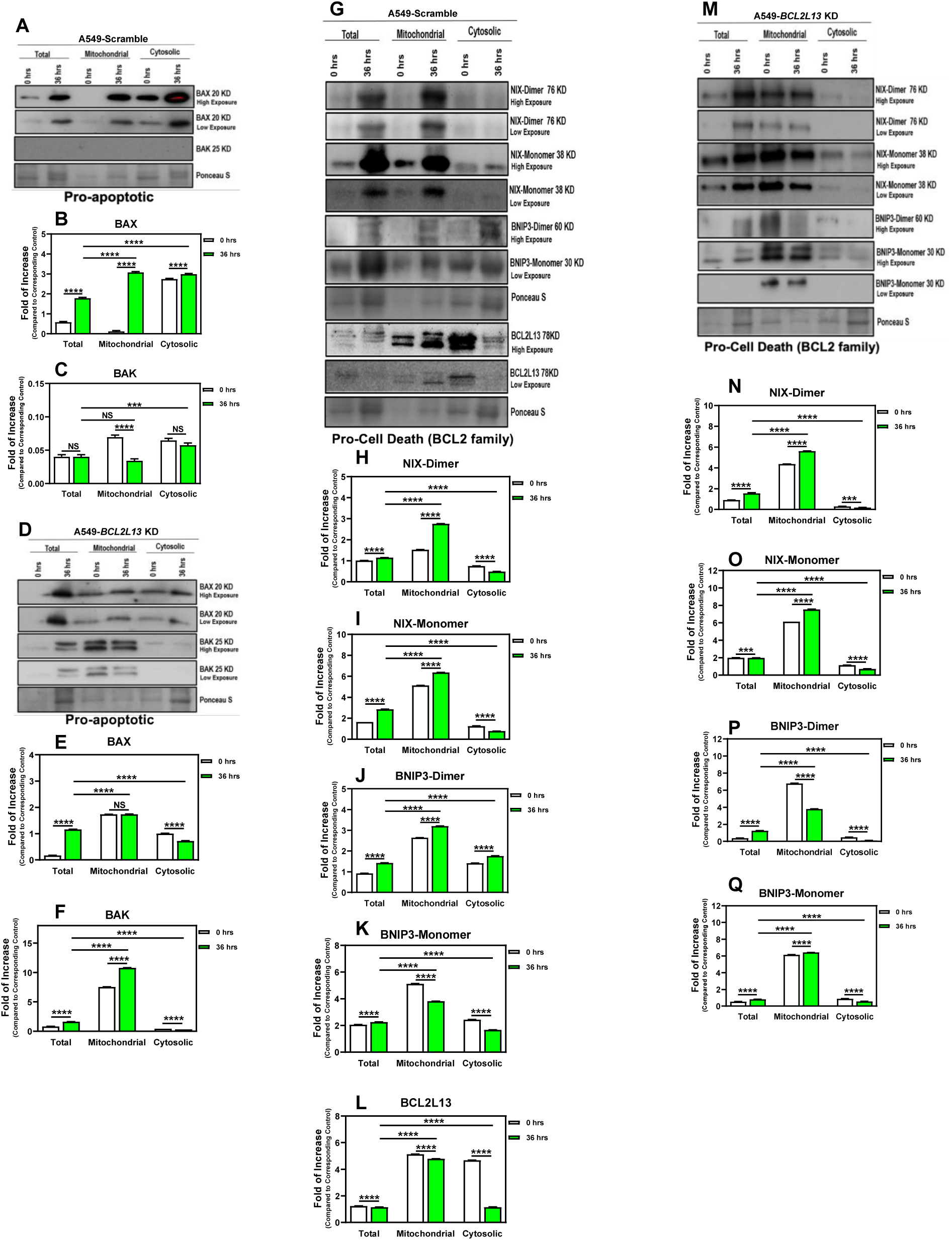
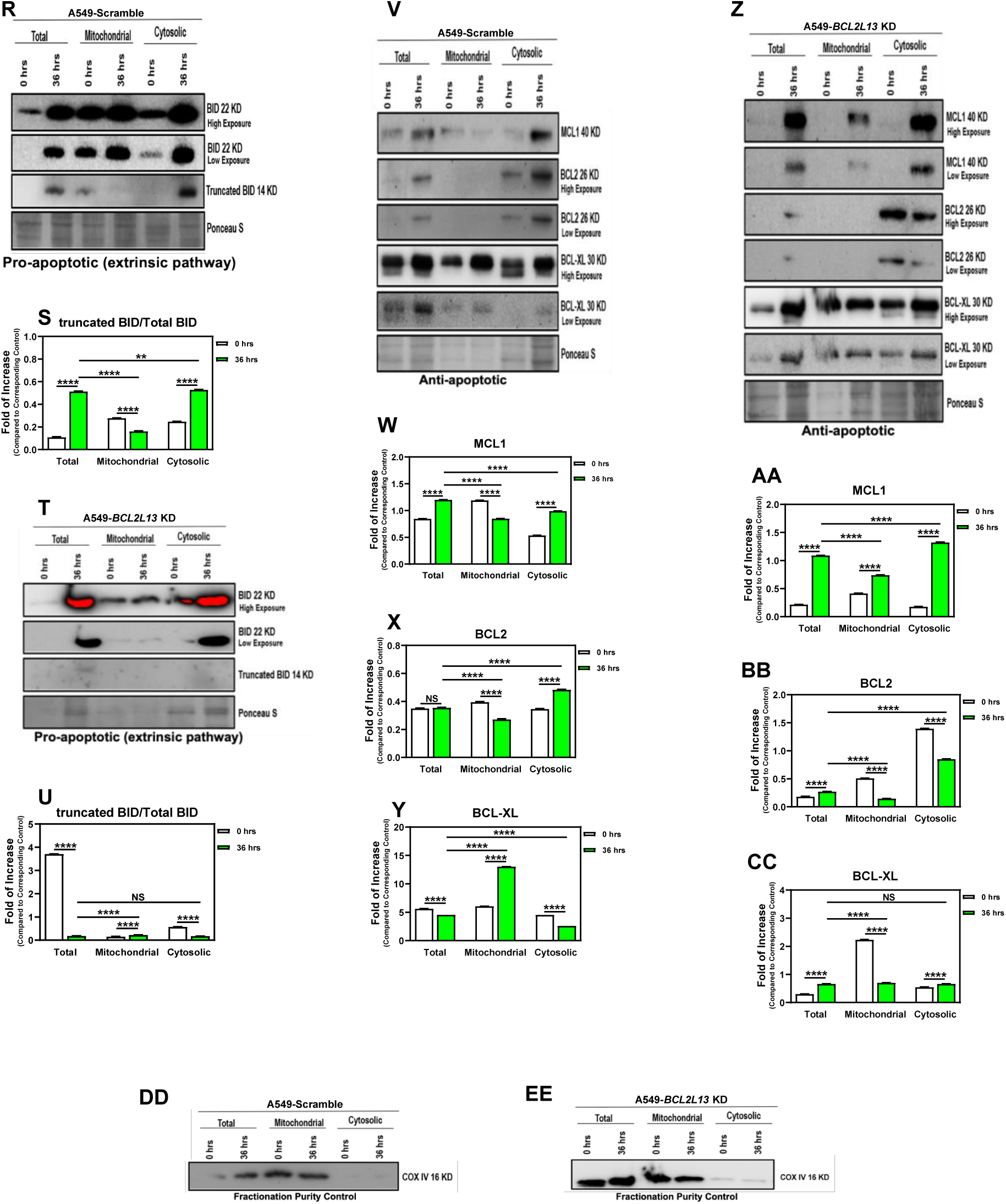
BCL2L13 depletion alters compartment-specific distribution of BCL2-family proteins during anoikis. Scramble and *BCL2L13* knockdown (KD) A549 cells were cultured under non-adherent conditions for 36 h, followed by mitochondrial and cytosolic fractionation and immunoblotting. Cells collected immediately after seeding on non-adherent plates served as the 0 h control. Protein abundance in mitochondrial and cytosolic fractions was normalized to total lysate. **(A–F)** BAX was enriched in mitochondrial fractions in both scramble and *BCL2L13*-KD cells, whereas BAK showed increased mitochondrial localization in *BCL2L13*-KD cells. **(G–Q)** NIX and BNIP3, including monomeric and dimeric forms, showed compartment-specific redistribution under anoikis conditions, with patterns differing between scramble and *BCL2L13*-KD cells. **(R–U)** BID processing showed a distinct pattern: in scramble cells, detachment increased the tBID/total BID signal in total and cytosolic fractions, whereas mitochondrial tBID was not increased. In *BCL2L13*-KD cells, tBID/total BID was markedly reduced after detachment, particularly in total and cytosolic fractions, with no clear mitochondrial enrichment. **(V–CC)** Anti-apoptotic BCL2-family proteins, including MCL1, BCL2 and BCL-XL, showed protein- and compartment-specific redistribution after detachment and BCL2L13 depletion. **(DD,EE)** COX IV was used to assess mitochondrial fraction enrichment, and Ponceau S staining was used to verify protein loading. Densitometric analysis was performed using AlphaEase FC and GraphPad Prism. Data are mean ± SEM from three independent experiments. NS, not significant; *p ≤ 0.05, **p ≤ 0.01, ***p ≤ 0.001, ****p ≤ 0.0001.

### Autophagy-marker changes accompany BCL2L13 depletion but do not explain anoikis resistance

Because autophagy can influence apoptosis and anoikis ^55^ ^56^, we asked whether altered autophagy contributes to the anoikis-resistant phenotype observed after *BCL2L13* knockdown. Pharmacological induction of autophagy with rapamycin or inhibition of late-stage autophagy with bafilomycin A1 did not significantly change viability in *BCL2L13*-KD, *BCL2L13*-OE or corresponding control A549 and LLC cells, as measured by MTT assay (Fig. 10A,B; Suppl. Fig. 5A,B). We next examined apoptosis under detachment conditions. PI Nicoletti analysis showed that neither rapamycin nor bafilomycin A1 significantly altered anoikis-associated apoptosis across the *BCL2L13*-KD, *BCL2L13*-OE or control groups (Fig. 10C,D; Suppl. Fig. 5C,D). Thus, acute pharmacological modulation of autophagy did not measurably rescue or enhance the anoikis phenotype in these models. Immunoblot analysis of detached A549 cells showed increased LC3-II accumulation together with enhanced p62 reduction in *BCL2L13*-KD cells compared with controls after 72 h of detachment (Fig. 10E–G). This marker profile is consistent with increased autophagy-associated turnover in *BCL2L13*-deficient cells. However, because pharmacological induction or inhibition of autophagy did not significantly alter viability or apoptosis under anoikis conditions, the increased autophagy-marker response does not appear to be sufficient to explain the detachment-survival advantage. Together, these data suggest that *BCL2L13* depletion is associated with increased autophagy-marker activity during detachment, but that *BCL2L13*-dependent anoikis resistance is not primarily driven by autophagy under the conditions tested. This supports a separation between the role of BCL2L13 in mitochondrial quality-control pathways and its effects on anchorage-independent survival.

**Figure 10.**
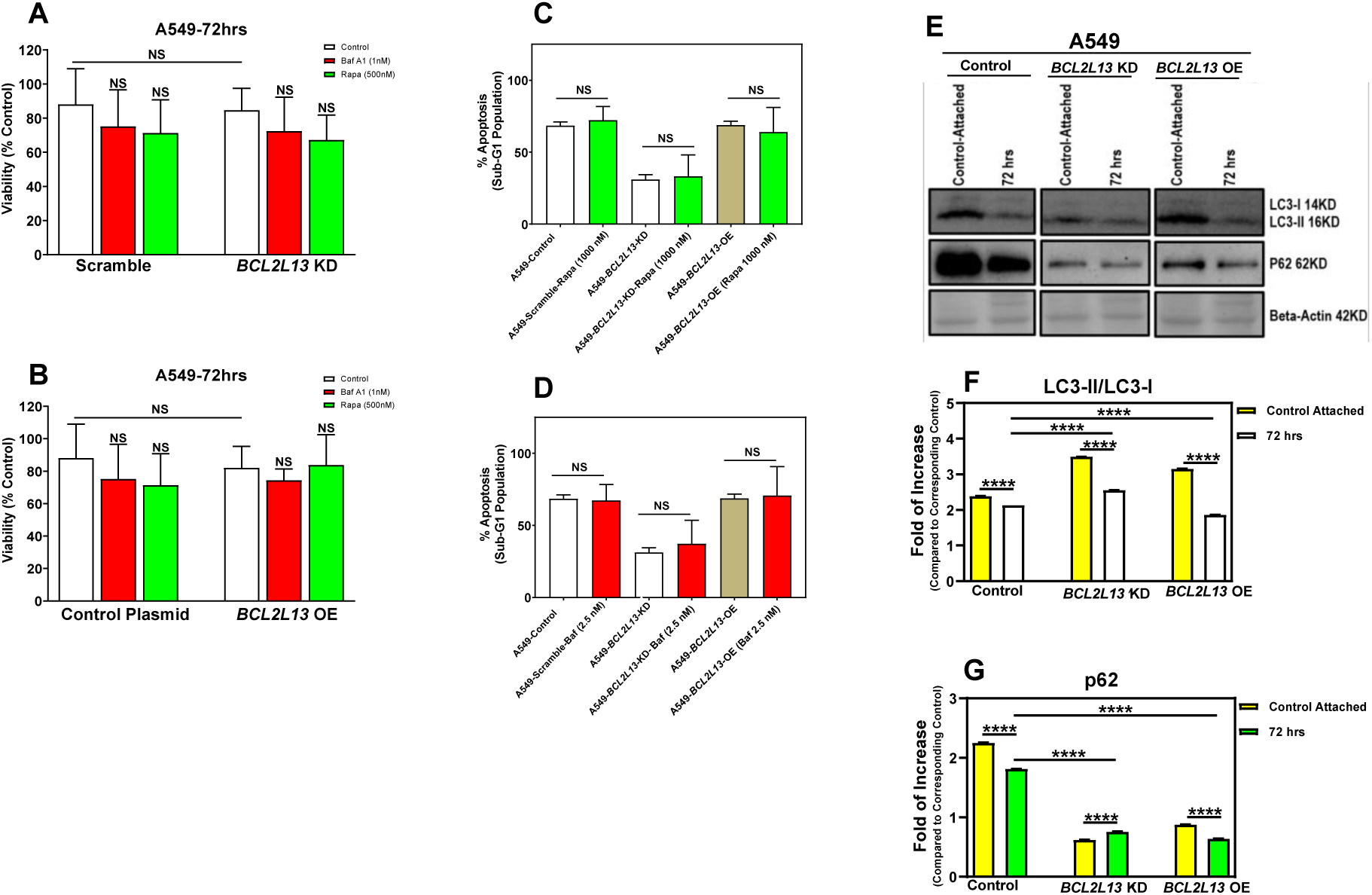
Autophagy modulation does not measurably alter BCL2L13-dependent anoikis responses in A549 cells. (A,B) *BCL2L13* knockdown (KD), *BCL2L13* overexpression (OE) and corresponding control A549 cells were cultured under non-adherent conditions and treated with bafilomycin A1 or rapamycin to inhibit late-stage autophagy or induce autophagy, respectively. Cell viability was assessed after 72 h by MTT assay and showed no significant treatment-dependent changes across the indicated groups. **(C,D)** Anoikis-associated apoptosis was assessed by PI Nicoletti assay after 72 h treatment with bafilomycin A1 or rapamycin. Pharmacological modulation of autophagy did not significantly alter apoptosis in control, *BCL2L13*-KD or *BCL2L13*-OE cells under detachment conditions. **(E–G)** Immunoblot analysis of autophagy markers under anoikis conditions showed increased LC3-II accumulation in *BCL2L13*-KD cells, whereas *BCL2L13*-OE reduced LC3-II levels relative to controls. p62/SQSTM1 abundance was quantified to assess autophagy-associated turnover. Attached cells were included as baseline controls; β-actin served as loading control. MTT data are mean ± SD from 15 measurements across three independent experiments; PI Nicoletti data are mean ± SD from three independent experiments with nine technical replicates; immunoblot quantification is mean ± SEM from three independent experiments. NS, not significant; *p ≤ 0.05, **p ≤ 0.01, ***p ≤ 0.001, ****p ≤ 0.0001.

### BCL2L13 expression is associated with detachment-dependent ceramide remodeling in NSCLC cells

Anoikis engages mitochondrial and membrane-associated death signaling, both of which can be influenced by ceramide metabolism. Ceramides have been implicated in mitochondrial apoptosis through effects on mitochondrial outer membrane permeabilization and cytochrome c release, and related mitochondrial apoptotic mechanisms have also been described in more recent studies ^8,57^. Ceramide channels in the outer mitochondrial membrane can be regulated by anti-apoptotic BCL2-family proteins such as BCL-XL ^58^, while ceramide-enriched membrane domains have been linked to BAX oligomerization ^59^. Ceramide composition may also affect adhesion-associated signaling, including Src localization and FAK activation ^60^. We therefore examined whether BCL2L13 expression was associated with altered ceramide-related lipid profiles under anoikis conditions.

Lipidomic profiling of detached A549 cells revealed a BCL2L13-dependent shift in ceramide-related species, including Cer, dhCer, MHC, DHC and THC classes (Fig. 11A). Under anoikis conditions, *BCL2L13*-OE cells showed higher abundance of several ceramide-related species, including Cer 24:0, Cer 24:1, dhCer 24:1, MHC 24:0, MHC 24:1, DHC 16:0 and DHC 24:0, whereas control and *BCL2L13*-KD cells showed comparatively lower abundance of these species (Fig. 11B–E). PCA separated control, KD and OE groups, indicating that *BCL2L13* expression state is associated with distinct lipidomic profiles during detachment (Fig. 11G). Volcano analysis further showed reduced abundance of several dhCer species in *BCL2L13*-KD cells, while no significant OE-associated species passed the same statistical thresholds (Fig. 11H,I). To identify lipid species contributing most strongly to group separation, we applied PLS-DA/VIP analysis and identified six candidate lipids with VIP scores >1.6 (Fig. 11J). Exploratory target prediction and interaction-network analysis of these lipid-associated features highlighted enrichment of mTOR-related pathways, with additional pathway and functional annotations linked to organelle-associated processes (Fig. 11K–N). These analyses should be interpreted as hypothesis-generating, because they do not directly measure signaling activity or protein–lipid interactions. Together, these data indicate that BCL2L13 expression is associated with context-dependent remodeling of ceramide-related lipids during anoikis. The lower C16- and C24-ceramide-associated profile in *BCL2L13*-KD cells compared with *BCL2L13*-OE cells differs from the pattern observed under adherent conditions, suggesting that detachment alters the relationship between BCL2L13 and sphingolipid metabolism. Given the reported links between C16/C24 ceramides, Src localization and FAK activation ^60^, these lipid changes may provide a plausible connection to the altered FAK phosphorylation observed in *BCL2L13*-deficient cells. However, direct biochemical validation will be required to determine whether ceramide remodeling functionally contributes to Src–FAK signaling or anoikis resistance in this model.

**Figure 11.**
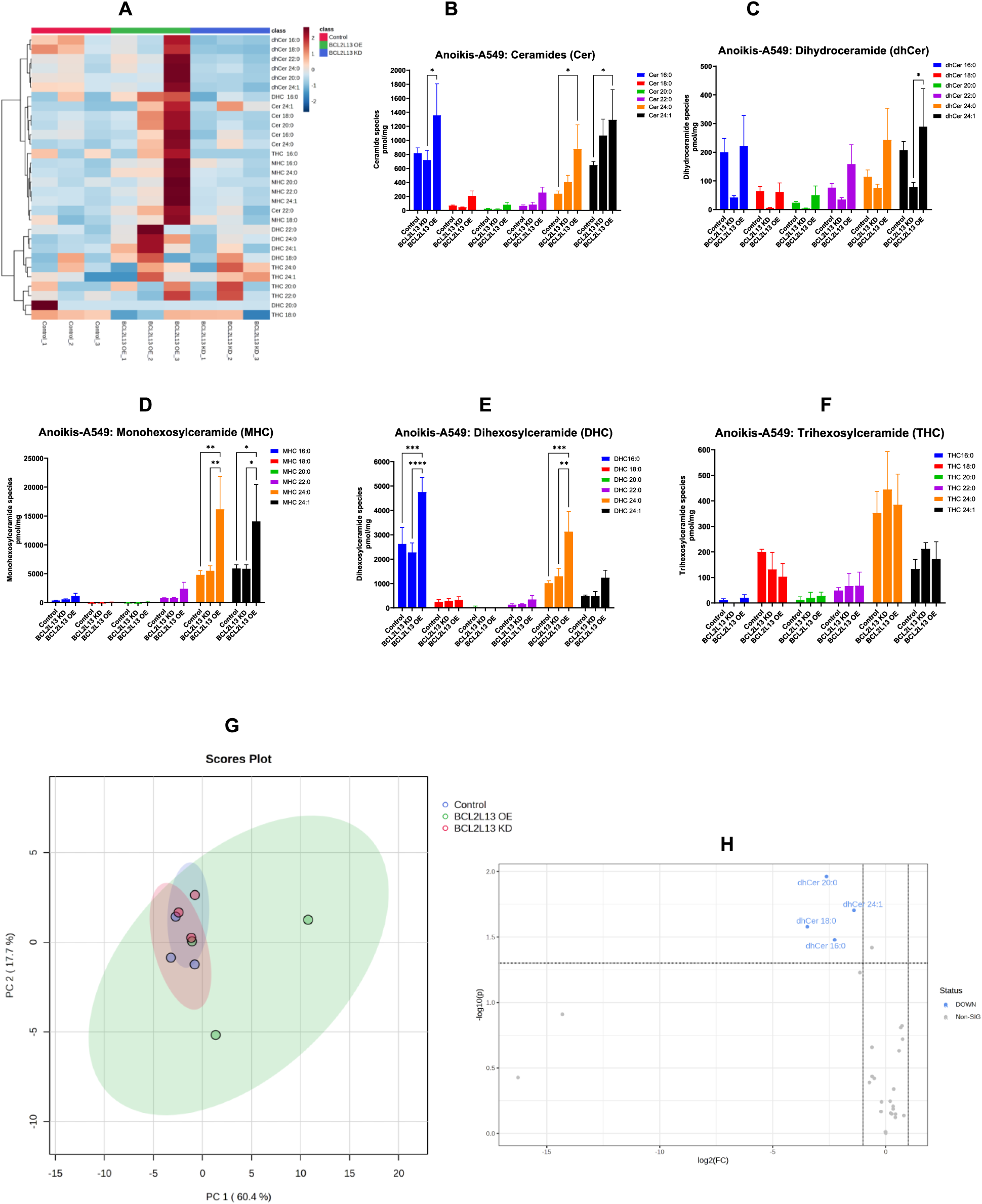
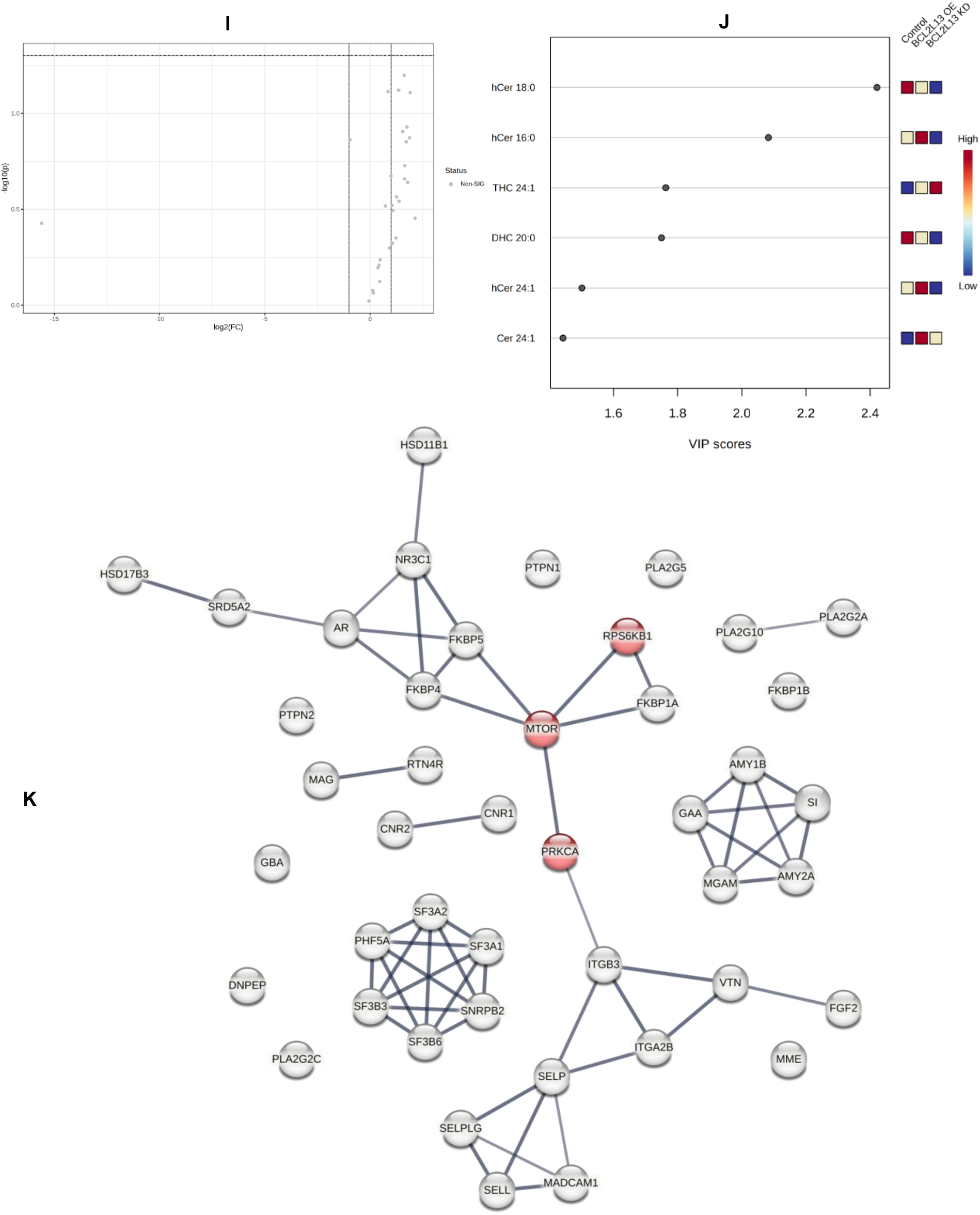
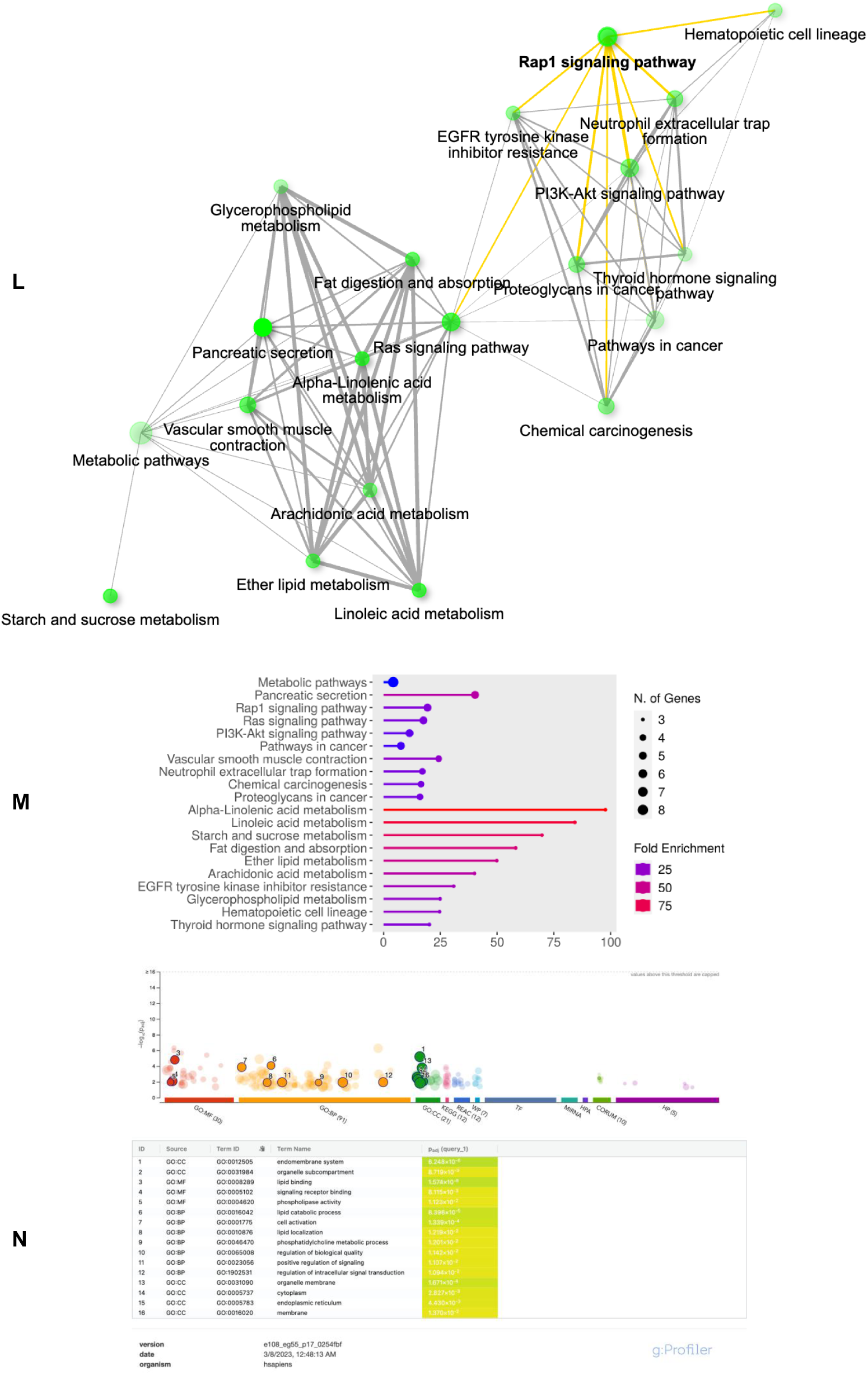
BCL2L13 expression state is associated with ceramide remodeling and mTOR-linked network annotations during anoikis. **(A)** Heatmap showing relative abundance of ceramide-related lipid species, including Cer, dhCer, MHC, DHC and THC, in A549 cells cultured under anoikis conditions. *BCL2L13* knockdown (KD), *BCL2L13* overexpression (OE) and their corresponding controls are shown. Scramble shRNA and empty-vector cells served as KD and OE controls, respectively. **(B–F)** Quantification of ceramide-family lipid species showing class- and species-selective changes after *BCL2L13*-KD or -OE. **(G)** Principal component analysis of ceramide-related lipid profiles from control, KD and OE cells under detachment conditions. **(H,I)** Volcano plots showing differentially abundant lipid species in *BCL2L13*-KD versus scramble control and *BCL2L13*-OE versus empty-vector control. **(J)** PLS-DA/VIP analysis identifying lipid species contributing to group separation. **(K)** Predicted protein-interaction network generated from targets associated with prioritized VIP lipids, highlighting mTOR-related pathway nodes. **(L,M)** KEGG pathway enrichment and pathway-interaction analysis of lipid-associated target networks. **(N)** Functional enrichment of molecular function, biological process and cellular component terms associated with the prioritized lipid-linked network. Network and enrichment analyses are exploratory and were used to annotate lipid-associated pathways rather than directly measure pathway activity. Data are mean ± SEM from three independent experiments. NS, not significant; *P ≤ 0.05, **P ≤ 0.01, ***P ≤ 0.001, ****P ≤ 0.0001 by two-way ANOVA.

### BCL2L13-dependent autophagy-marker responses differ between attached and anoikis states

Lipidomic network analyses highlighted PI3K–AKT–mTOR-related pathways among the BCL2L13-associated signatures identified under both attached and anoikis conditions. Because PI3K–AKT–mTOR signalling is a major upstream regulator of autophagy ^61^, and mitophagy depends on core autophagy machinery linked to this pathway ^62^, we asked whether the experimental autophagy-marker data were consistent with these pathway-level annotations. Under attached conditions, *BCL2L13* knockdown reduced TGFβ1-associated mitophagy marker responses, including lower LC3B-II/LC3B-I ratio and reduced p62 turnover in cytosolic fractions (Fig. 3), consistent with a role for BCL2L13 in mitophagy-associated processing in adherent NSCLC cells. This interpretation is in line with previous evidence connecting PI3K–AKT–mTOR signalling to autophagy and mitophagy regulation (Chen, Zhang et al. 2022). By contrast, under anoikis conditions, *BCL2L13*-KD cells showed increased LC3B-II/LC3B-I ratio together with increased p62 reduction (Fig. 10), indicating a different autophagy-marker pattern during detachment. Thus, *BCL2L13* loss was associated with reduced mitophagy-associated responses in attached cells but increased autophagy-marker turnover under anoikis conditions. Together, these findings suggest that BCL2L13 influences autophagy-related responses in a context-dependent manner. The lipidomics-based PI3K–AKT–mTOR enrichment provides a plausible pathway-level link, but these data should be interpreted as supportive rather than direct evidence of pathway activation. Direct measurement of PI3K–AKT–mTOR activity will be required to define how this signalling axis contributes to BCL2L13-dependent autophagy and survival responses in NSCLC cells.

## Discussion

This study identifies BCL2L13 as a context-dependent regulator of mitochondrial quality control, epithelial–mesenchymal plasticity and detachment survival in NSCLC adenocarcinoma cells. Across human tissue analysis and mechanistic cell models, BCL2L13 showed compartment- and subtype-dependent expression, was required for efficient mitophagy-associated responses, and influenced anoikis sensitivity, mitochondrial function and sphingolipid-state remodeling. Rather than acting through a single linear pathway, BCL2L13 appears to operate at the intersection of mitochondrial turnover, BCL2-family signaling, adhesion-associated survival pathways and lipid metabolism.

The human TMA and matched patient samples provided an important translational frame for the experimental work. BCL2L13 immunoreactivity was predominantly cytoplasmic/granular, consistent with mitochondrial-associated localization, and was more evident in NSCLC components than in small cell carcinoma. In adenocarcinoma and squamous cell carcinoma, BCL2L13 staining was retained in confirmed lymph-node metastases in selected cases, although nodal staining was heterogeneous and dependent on the presence of viable metastatic tumor. These findings do not establish BCL2L13 as a prognostic biomarker, but they support its biological relevance in human lung cancer and justify mechanistic analysis in NSCLC models.

BCL2L13 has previously been described as a mitophagy receptor, but its role in NSCLC had remained unclear ^40,63–65^. In both A549 and LLC cells, BCL2L13 depletion reduced TGFβ1-associated LC3β–mitochondria colocalization, TOMM20–LAMP1 overlap, mitochondrial LC3B-II accumulation and TOMM20/p62 turnover. These effects were not compensated by BNIP3 or NIX, supporting a non-redundant role for BCL2L13 in this setting ^66^. CCCP experiments further supported this interpretation: depolarization-induced mitophagy-associated trafficking was evident in control cells but attenuated after *BCL2L13* knockdown. Thus, BCL2L13 is not simply redistributed during mitochondrial stress; it is required for efficient mitophagy-associated processing under both TGFβ1- and CCCP-driven conditions.

This mitochondrial phenotype was closely linked to EMT-associated plasticity. TGFβ1 is a well-established inducer of EMT in NSCLC, and previous work has connected autophagy to EMT regulation in this system ^11^. Here, *BCL2L13* knockdown enhanced TGFβ1-associated mesenchymal marker expression and increased migration, whereas *BCL2L13* overexpression partially preserved epithelial features. Importantly, CCCP reduced EMT-associated markers in control cells, but this effect was not reproduced in *BCL2L13*-deficient cells. These results support a model in which effective mitochondrial quality control restrains EMT-associated phenotypes, while *BCL2L13* loss permits mitochondrial stress to coexist with increased plasticity. This interpretation is consistent with LC3-dependent BCL2L13 activity and the requirement for intact autophagy machinery in mitophagy ^57,63^, but it does not imply that bulk autophagy alone explains the phenotype ^67^.

BCL2L13 also influenced mitochondrial function, although the response was not a simple gain– loss relationship. Both knockdown and overexpression reduced several OCR-derived parameters, while *BCL2L13* knockdown increased proton leak and *BCL2L13* overexpression increased mitochondrial ROS. These findings suggest that *BCL2L13* expression must be balanced for mitochondrial respiratory homeostasis. The decrease in respiratory capacity in *BCL2L13*-deficient cells is compatible with metabolic adaptation often observed in cancer models ^68–71^, but the current data do not demonstrate a full metabolic switch. Increased TMRM signal in both knockdown and overexpression cells indicate altered membrane-potential regulation rather than a uniform oxidative-stress phenotype ^72,73^.

A second major finding is that *BCL2L13* depletion reduced detachment-induced apoptosis. Anoikis resistance is central to metastatic competence, and *BCL2L13* loss increased survival under anchorage-independent conditions in both A549 and LLC cells. However, this phenotype was not explained by a proportional change in canonical caspase activity. In A549 cells, *BCL2L13* knockdown reduced caspase-8 and caspase-3/7 activity, while overexpression increased PI-defined apoptosis but reduced caspase activity. This non-parallel relationship suggests that BCL2L13 affects detachment survival through mitochondrial and adhesion-linked regulatory states rather than a single caspase-dependent mechanism. Non-caspase mitochondrial effectors, including AIF or EndoG^74^., remain plausible contributors but were not directly tested

Analysis of BCL2-family proteins supported this more complex interpretation. *BCL2L13* depletion did not globally suppress pro-apoptotic proteins: BAK, BNIP3 and NIX were increased or redistributed, whereas BAX was not uniformly reduced. Because BAX and BAK cooperate to execute mitochondrial outer membrane permeabilization, altered balance or activation state may be more important than total abundance alone ^75–77^. At the same time, increased MCL1 and BCL2 may contribute to a survival-biased state ^78,79^. BNIP3 and NIX, although increased in some anoikis conditions, did not restore apoptosis, reinforcing the idea that BCL2L13 has a non-redundant role in coordinating mitochondrial fate decisions (Pedanou, Gobeil et al. 2016).

Subcellular fractionation refined this model. *BCL2L13* depletion did not prevent mitochondrial recruitment of BAX, BAK, BNIP3 or NIX during detachment. Instead, BID processing showed a distinct pattern: in control cells, detachment increased tBID/total BID mainly in total and cytosolic fractions, while mitochondrial tBID was not increased; in *BCL2L13*-deficient cells, tBID/total BID was markedly reduced after detachment with no clear mitochondrial enrichment. Thus, anoikis resistance in *BCL2L13*-deficient cells is unlikely to arise simply from failed mitochondrial recruitment of pro-apoptotic proteins. A more likely interpretation is that *BCL2L13* loss alters the coupling between apoptotic priming, BID processing and mitochondrial execution.

FAK signaling provided an additional layer of detachment-associated regulation. *BCL2L13* knockdown reduced the p-FAK Tyr576/577/FAK ratio while increasing p-FAK Tyr925/FAK, suggesting altered signaling quality rather than uniform FAK suppression or activation. Tyr576/577 is linked to catalytic activation downstream of Src-dependent adhesion signaling, whereas Tyr925 has been associated with FAK–Src/MAPK signaling ^80–82^. This pattern is consistent with a shift from canonical adhesion-linked signaling toward alternative survival-associated signaling in detached cells, but causal experiments targeting FAK or Src are needed before assigning mechanism.

The lipidomics data add a mechanistically important, but still hypothesis-generating, dimension. *BCL2L13* has been reported to interact with CerS2 and CerS6^10^, and ceramide metabolism has established links to mitochondrial apoptosis, BAX activation and cell-death regulation ^83^. Under attached conditions, *BCL2L13* knockdown selectively increased several C24-containing sphingolipid species, including dhCer 24:0, MHC 24:0 and THC 24:0. This is consistent with altered CerS2/6-associated lipid metabolism ^84–86^, but does not directly demonstrate altered CerS activity or BCL2L13–CerS complex formation. Therefore, the lipidomic data should be interpreted as evidence of BCL2L13-associated sphingolipid remodeling, not as proof of direct enzymatic regulation.

Detachment changed this relationship. Under anoikis conditions, *BCL2L13* overexpression was associated with increased ceramide-related species, whereas *BCL2L13* knockdown showed comparatively lower C16- and C24-associated ceramide profiles. This inversion suggests that adhesion state modifies the relationship between BCL2L13 and sphingolipid metabolism. Given that ceramides can promote mitochondrial apoptosis ^87^ and that integrin–FAK signaling regulates anoikis ^88,89^, the detachment-dependent ceramide profile offers a plausible lipid-based explanation for altered survival signaling. However, this connection remains inferential. Direct assessment of Src recruitment, FAK activity, ceramide localization and CerS2/6 function will be required to determine whether ceramide remodeling is upstream of the anoikis phenotype.

Ceramides may also link BCL2L13 to mitochondrial quality control. Mitochondrial ceramide can recruit LC3B-II-positive autophagic membranes and promote mitophagy ^36,90^, while ceramide accumulation has been linked to impaired mitochondrial respiration ^91^. In this context, BCL2L13 may integrate lipid state with selective mitochondrial turnover ^63,64^. The pathway-annotation analyses further highlighted PI3K–AKT and mTOR-related networks, which are biologically plausible given their roles in survival and autophagy. However, these analyses were exploratory and do not directly measure pathway activation. They are best viewed as a framework for prioritizing future biochemical validation.

A key insight from the study is that BCL2L13 has different effects in attached versus detached states. In adherent cells, *BCL2L13* loss impaired TGFβ1-associated mitophagy and enhanced EMT-associated phenotypes. In detached cells, the same perturbation reduced apoptosis, altered BID processing, changed FAK phosphorylation and reshaped ceramide-related lipid profiles. Pharmacological modulation of autophagy did not measurably affect viability or apoptosis under anoikis conditions, despite increased LC3-II and p62 reduction in *BCL2L13*-deficient cells. These results separate the role of BCL2L13 in mitochondrial quality control from its role in anchorage-independent survival, and suggest that detachment rewires the downstream consequences of BCL2L13-dependent mitochondrial and lipid signaling.

The innovation of this work lies in connecting BCL2L13-dependent mitophagy to metastatic traits through two experimentally distinct states: adherent EMT induction and anchorage-independent survival. The human tissue data position BCL2L13 within clinically relevant lung cancer compartments, while the cell-based experiments link BCL2L13 to mitophagy, mitochondrial bioenergetics, BCL2-family redistribution, FAK phosphorylation and sphingolipid remodeling. This integrated framework provides a translational rationale for studying BCL2L13 not as a single-pathway death regulator, but as a mitochondrial–lipid signaling node whose consequences depend on adhesion state and tumor context.

The study is limited by the absence of rescue experiments, direct CerS2/6 activity measurements, organelle-resolved lipidomics and direct tests of Src–FAK or PI3K–AKT–mTOR signaling. Whole-cell lipidomics cannot define mitochondrial ceramide pools, and TMA findings require validation in larger, independently annotated cohorts. Future studies should restore BCL2L13 in knockdown cells, map BCL2L13–CerS2/6 complexes under attached and anoikis conditions, quantify mitochondrial sphingolipid species, and test whether FAK/Src or mTOR modulation reverses detachment survival ^92–99^.

Taken together, our findings support a model in which BCL2L13 coordinates mitochondrial quality control, lipid-state remodeling and detachment-associated survival in NSCLC adenocarcinoma cells. Its function is not fixed but context-dependent: in adherent cells, BCL2L13 supports mitophagy and limits EMT-associated plasticity, whereas during detachment its loss favors anoikis resistance through altered apoptotic coupling, FAK signaling and ceramide-associated lipid states. This places BCL2L13 at a mechanistic interface between mitochondrial homeostasis and metastatic adaptation in NSCLC.

## Author contributions

J.A. designed and performed the main experimental work, including subcellular fractionation, immunoblotting, transfection, fluorescence microscopy, ImageJ-based image analysis, data analysis and interpretation, and drafted the manuscript. S.C.d.S.R. and R.V. contributed to lipidomics data analysis, interpretation, visualization, and figure preparation. A.S. performed Seahorse extracellular flux assays, analyzed mitochondrial bioenergetics data, and prepared the corresponding figures. S.D mitochondrial biogenesis M.A., Z.B performed correlation analyses and statistical assessment of tissue microarray data. A.G. performed real-time cell migration assays, analyzed migration data, and generated the corresponding figures. A.R. provided the lipidomics platform, technical support and expertise for lipidomics studies, and contributed to lipidomics interpretation. S.H.-K. evaluated and interpreted immunohistochemistry data and provided pathology-related feedback. M.M. and A.BB performed data-mining analyses and generated associated figures. J.W.G. BCL2L13 over expression study B.K. and N.A. contributed scientific input, clinical data interpretation, and critical revision of the manuscript. S.G. conceived and supervised the study, contributed to experimental design, data interpretation, manuscript writing and revision, and coordinated the overall project. All authors reviewed and approved the final version of the manuscript.

## Appendix

**Supplementary Figure 1.**
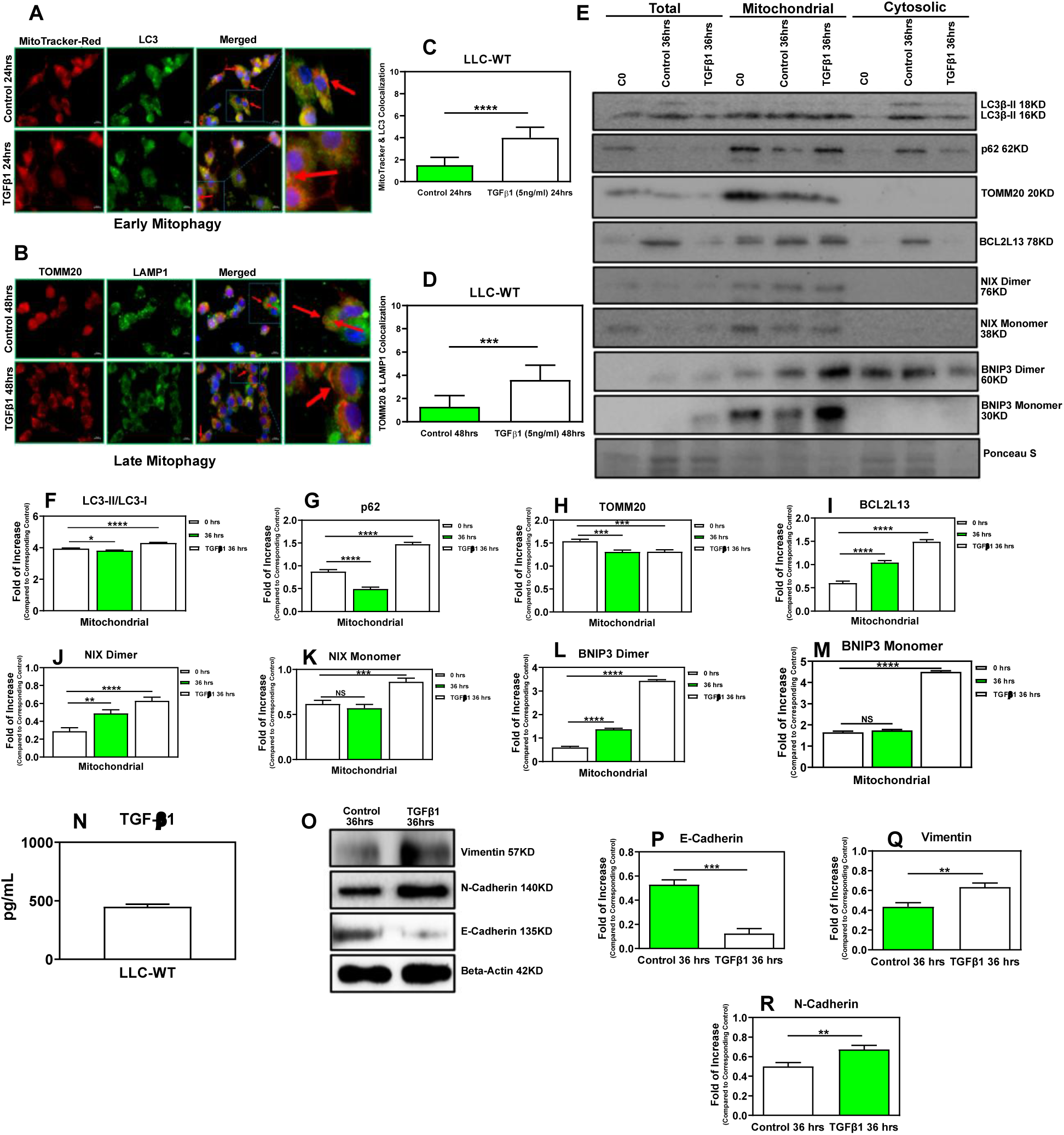
TGFβ1 induces mitophagy-associated responses and EMT marker changes in LLC cells. **(A–D)** LLC cells treated with TGFβ1 showed increased LC3β–mitochondria colocalization at 24 h and increased TOMM20–LAMP1 overlap at 48 h, consistent with early and late mitophagy-associated trafficking. **(E–M)** Subcellular fractionation and immunoblotting at 36 h showed increased mitochondrial LC3-II, altered p62 abundance and reduced TOMM20, together with mitochondrial enrichment of BCL2L13 and BNIP3; NIX showed a weaker response. Ponceau S was used to assess loading/fraction quality. **(N)** ELISA confirmed active TGFβ1 secretion at 36 h. **(O–R)** TGFβ1 treatment induced EMT-associated marker changes, including reduced E-cadherin and increased vimentin and N-cadherin. β-actin served as loading control for whole-cell immunoblots. Cells were treated with TGFβ1, 5 ng/ml, where indicated. Scale bars, 10 μm. Data are mean ± SEM from three independent experiments; imaging quantification was performed from 10 fields with approximately five cells per field. NS, not significant; *p ≤ 0.05, **p ≤ 0.01, ***p ≤ 0.001, ****p ≤ 0.0001.

**Supplementary Figure 2.**
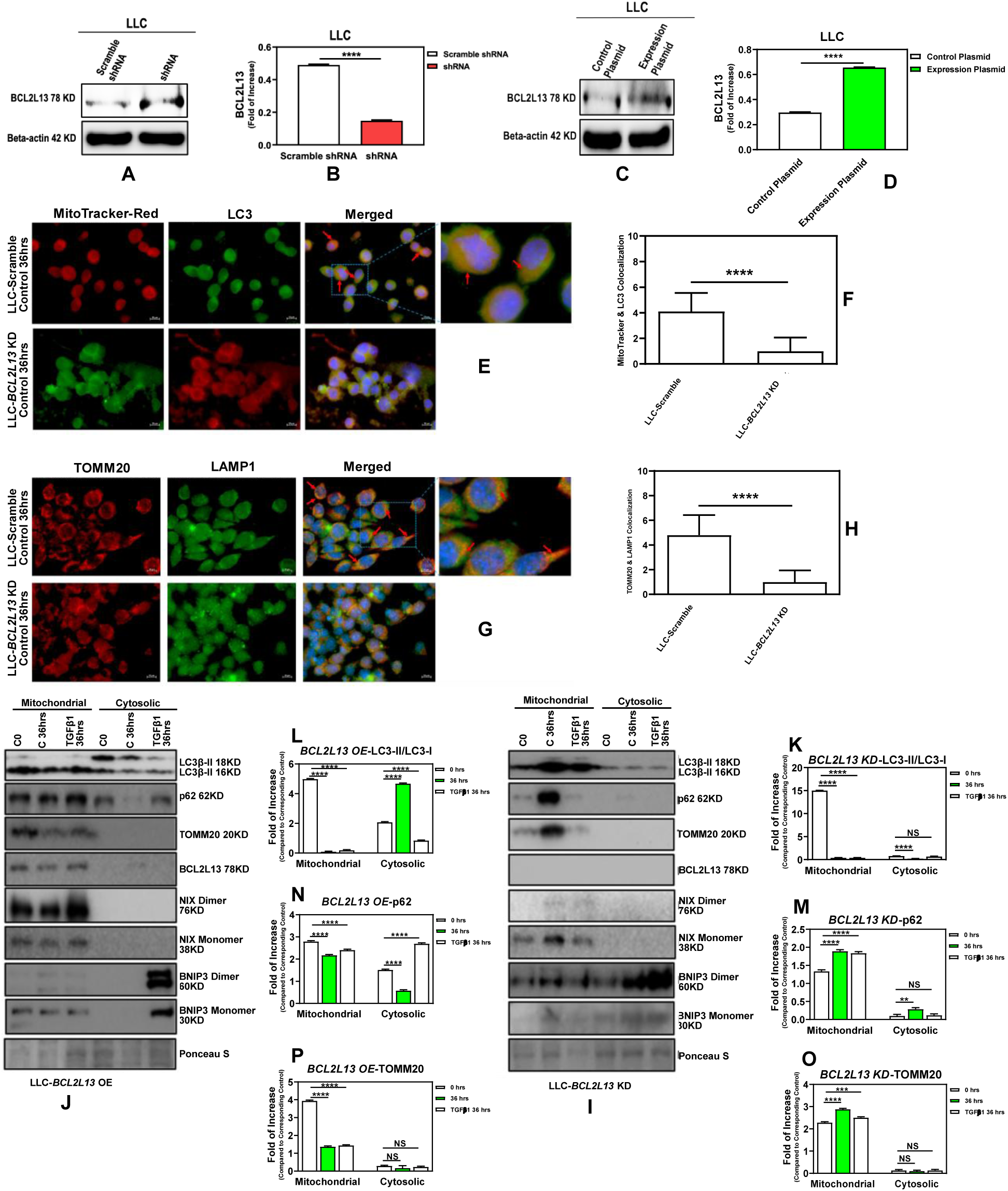
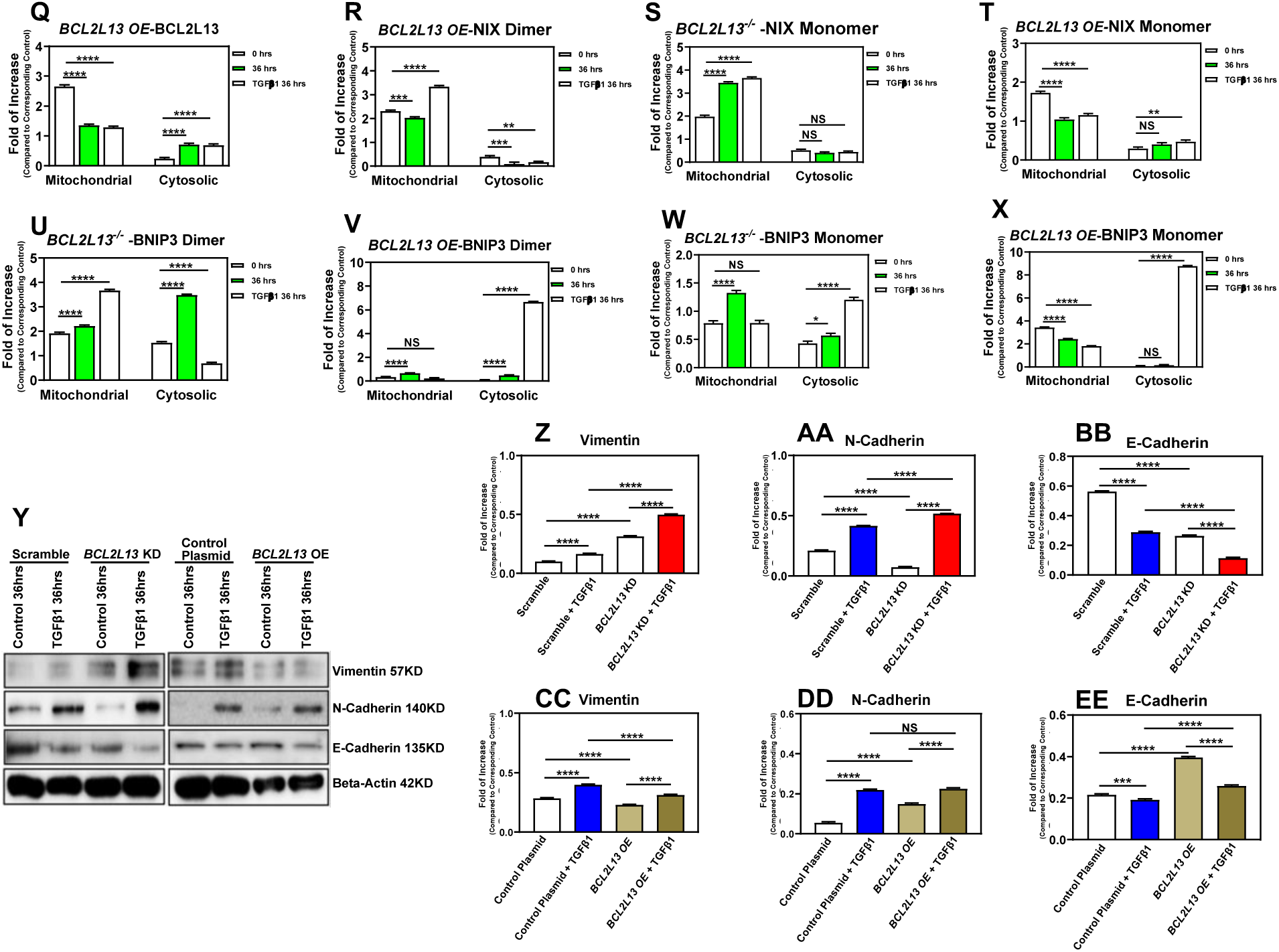
*Bcl2l13* depletion reduces mitophagy-associated responses and enhances EMT marker changes in LLC cells. **(A–D)** Stable *Bcl2l13* knockdown (KD) and overexpression (OE) were generated in LLC cells and validated by immunoblotting. Scramble shRNA and empty vector were used as the respective controls; β-actin served as loading control. **(E–H)** Immunocytochemical analysis after TGFβ1 treatment showed reduced LC3β–mitochondria colocalization and TOMM20–LAMP1 overlap in *Bcl2l13*-KD cells, consistent with attenuated mitophagy-associated trafficking. **(I–P)** Subcellular fractionation and immunoblotting showed reduced mitochondrial LC3-II accumulation and altered p62 and TOMM20 turnover after *Bcl2l13* depletion, whereas *Bcl2l13*-OE showed increased mitophagy-associated cargo processing. (Q–X) *Bcl2l13* perturbation was associated with changes in the mitochondrial and cytosolic distribution of NIX and BNIP3 isoforms. **(Y–EE)** EMT-marker analysis showed that *Bcl2l13*-KD enhanced TGFβ1-associated vimentin induction and E-cadherin loss, whereas *Bcl2l13*-OE partially opposed these changes. Cells were treated with TGFβ1, 5 ng/ml, for 36 h where indicated. Scale bars, 10 μm. Data are mean ± SEM from three independent experiments; imaging quantification was performed from 10 fields with approximately five cells per field. NS, not significant; *p ≤ 0.05, **p ≤ 0.01, ***p ≤ 0.001, ****p ≤ 0.0001.

**Supplementary Figure 3.**
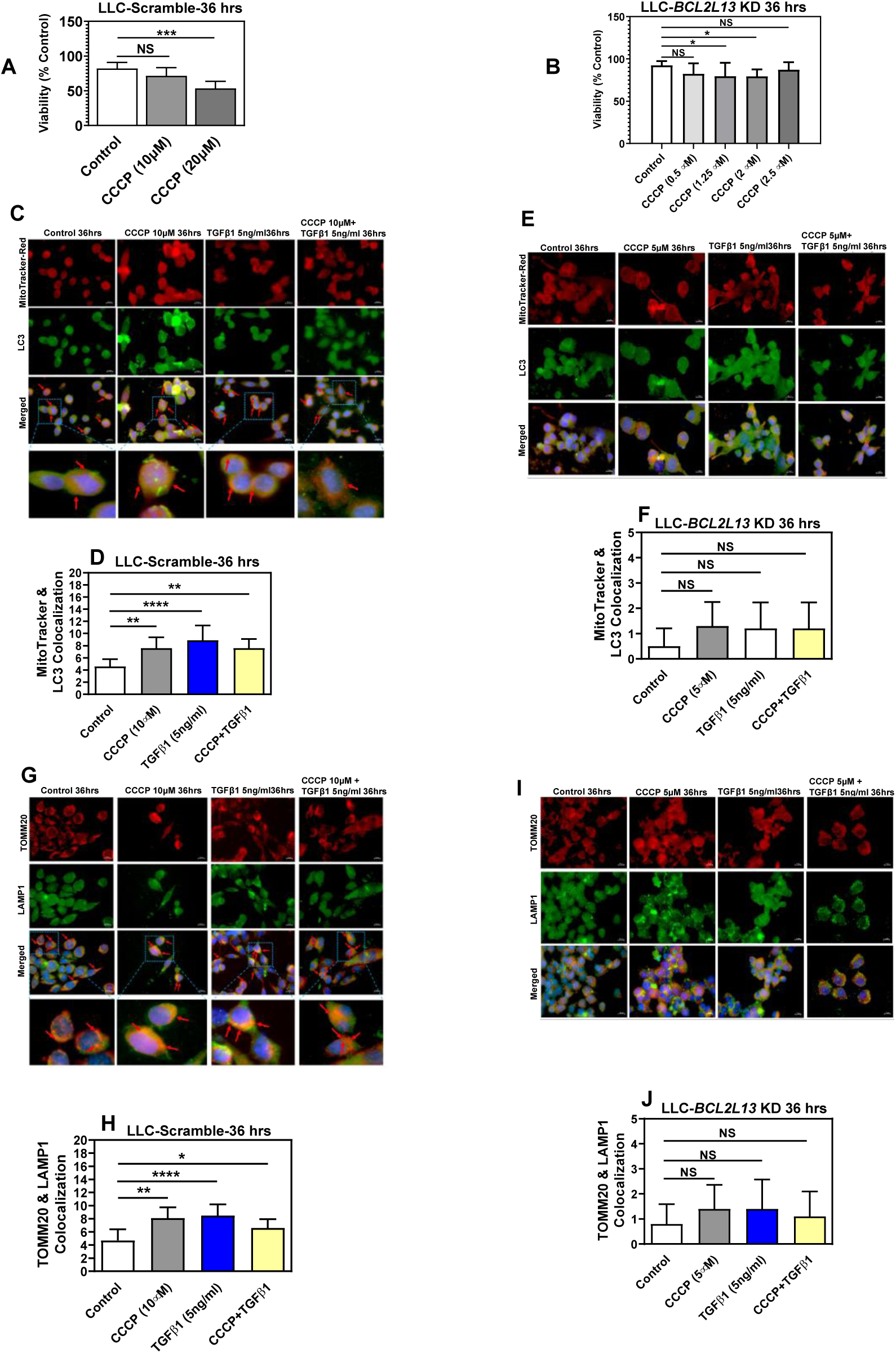
*Bcl2l13* depletion attenuates CCCP-associated mitophagy responses in LLC cells. **(A,B)** MTT-based dose optimization of CCCP in scramble and *Bcl2l13* knockdown (KD) LLC cells after 48 h treatment with 2.5–20 µM CCCP. DMSO-treated cells served as controls. **(C–F)** Immunocytochemical analysis after CCCP, TGFβ1, or combined CCCP/TGFβ1 treatment showed increased LC3β–mitochondria colocalization in scramble cells, whereas this response was not significantly induced in *Bcl2l13*-KD cells. **(G–J)** TOMM20–LAMP1 colocalization showed a similar pattern, with increased overlap in scramble cells but reduced mitophagy-associated trafficking in *Bcl2l13*-KD cells. Cells were treated with TGFβ1, 5 ng/ml, where indicated; ICC analyses were performed after 36 h treatment. Scale bars, 10 μm. MTT data are mean ± SD from 15 measurements across three independent experiments; ICC quantification is mean ± SEM from three independent experiments. NS, not significant; *p ≤ 0.05, **p ≤ 0.01, ***p ≤ 0.001, ****p ≤ 0.0001.

**Supplementary Figure 4.**
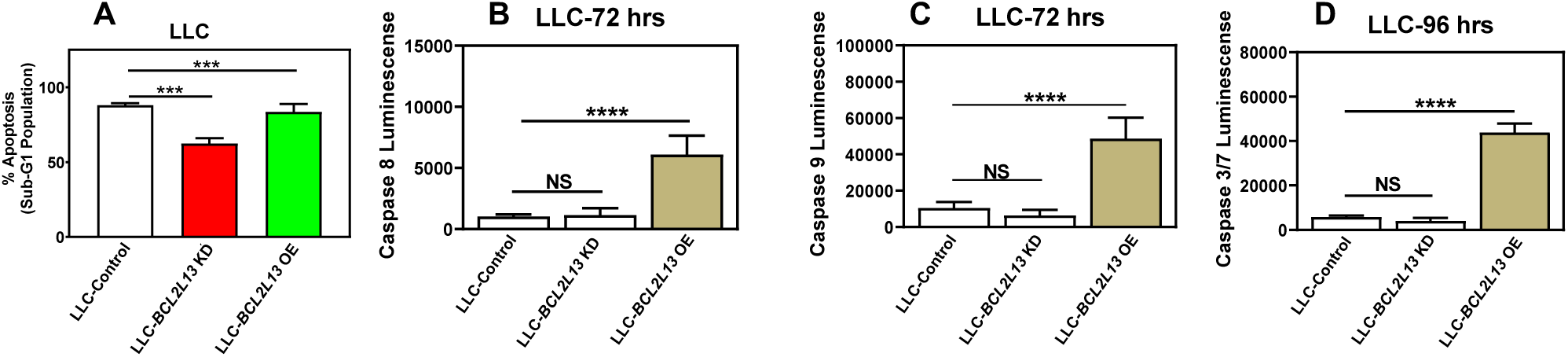
*Bcl2l13* depletion reduces anoikis-associated apoptosis in LLC cells with limited caspase changes. **(A)** Anoikis-associated apoptosis was assessed in *Bcl2l13* knockdown (KD), *Bcl2l13* overexpression (OE) and corresponding control LLC cells after 96 h culture under non-adherent conditions using the PI Nicoletti assay. *Bcl2l13*-KD reduced detachment-induced apoptosis relative to scramble control, whereas *Bcl2l13*-OE increased apoptosis relative to empty-vector control. **(B–D)** Caspase-Glo assays showed that *Bcl2l13*-KD did not significantly alter caspase-8, caspase-9 or caspase-3/7 activity under anoikis conditions, while *Bcl2l13*-OE increased activity of all three caspases. Caspase-8 and caspase-9 were measured at 72 h, and caspase-3/7 at 96 h. PI Nicoletti data are mean ± SD from three independent experiments with nine technical replicates; caspase data are mean ± SEM from three independent experiments. NS, not significant; *p ≤ 0.05, **p ≤ 0.01, ***p ≤ 0.001, ****p ≤ 0.0001.

**Supplementary Figure 5.**
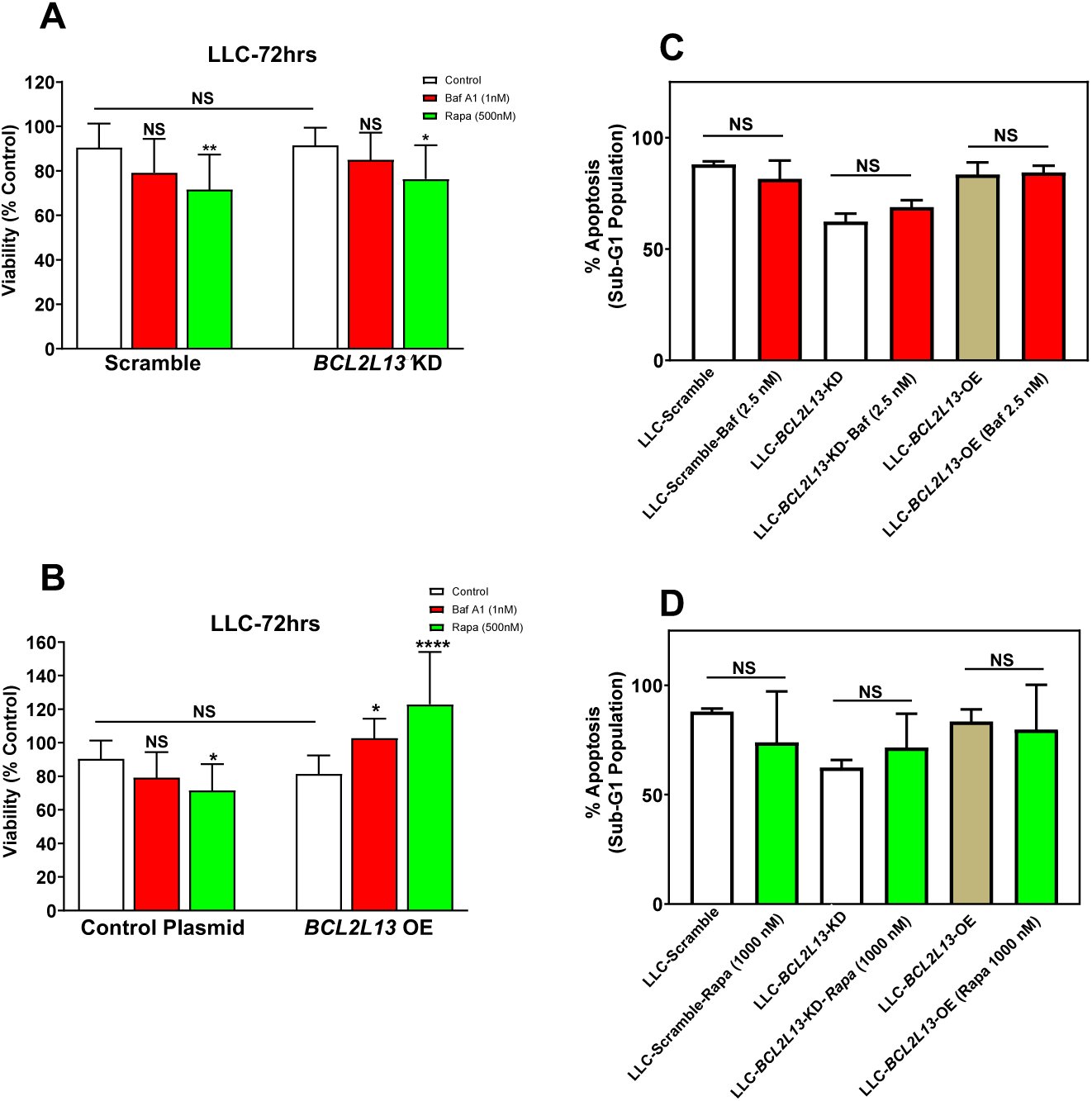
Autophagy modulation has limited effect on anoikis-associated apoptosis in LLC cells. **(A,B)** *Bcl2l13* knockdown (KD), *Bcl2l13* overexpression (OE) and corresponding control LLC cells were cultured under non-adherent conditions and treated with bafilomycin A1 or rapamycin to inhibit late-stage autophagy or induce autophagy, respectively. Cell viability was assessed after 72 h by MTT assay. Rapamycin reduced viability in control and *Bcl2l13*-KD cells but increased viability in *Bcl2l13*-OE cells, indicating an expression-state-dependent viability response. **(C,D)** Anoikis-associated apoptosis was assessed by PI Nicoletti assay after 72 h treatment with bafilomycin A1 or rapamycin. Autophagy modulation did not significantly alter apoptosis across control, *Bcl2l13*-KD or *Bcl2l13*-OE groups under detachment conditions. For MTT assays, cells were treated with bafilomycin A1, 1 nM, or rapamycin, 500 nM; for PI Nicoletti assays, cells were treated with bafilomycin A1, 2.5 nM, or rapamycin, 1000 nM. MTT data are mean ± SD from 15 measurements across three independent experiments; PI Nicoletti data are mean ± SD from three independent experiments with nine technical replicates. NS, not significant; *p ≤ 0.05, **p ≤ 0.01, ***p ≤ 0.001, ****p ≤ 0.0001.

**Supplementary Table 1.**
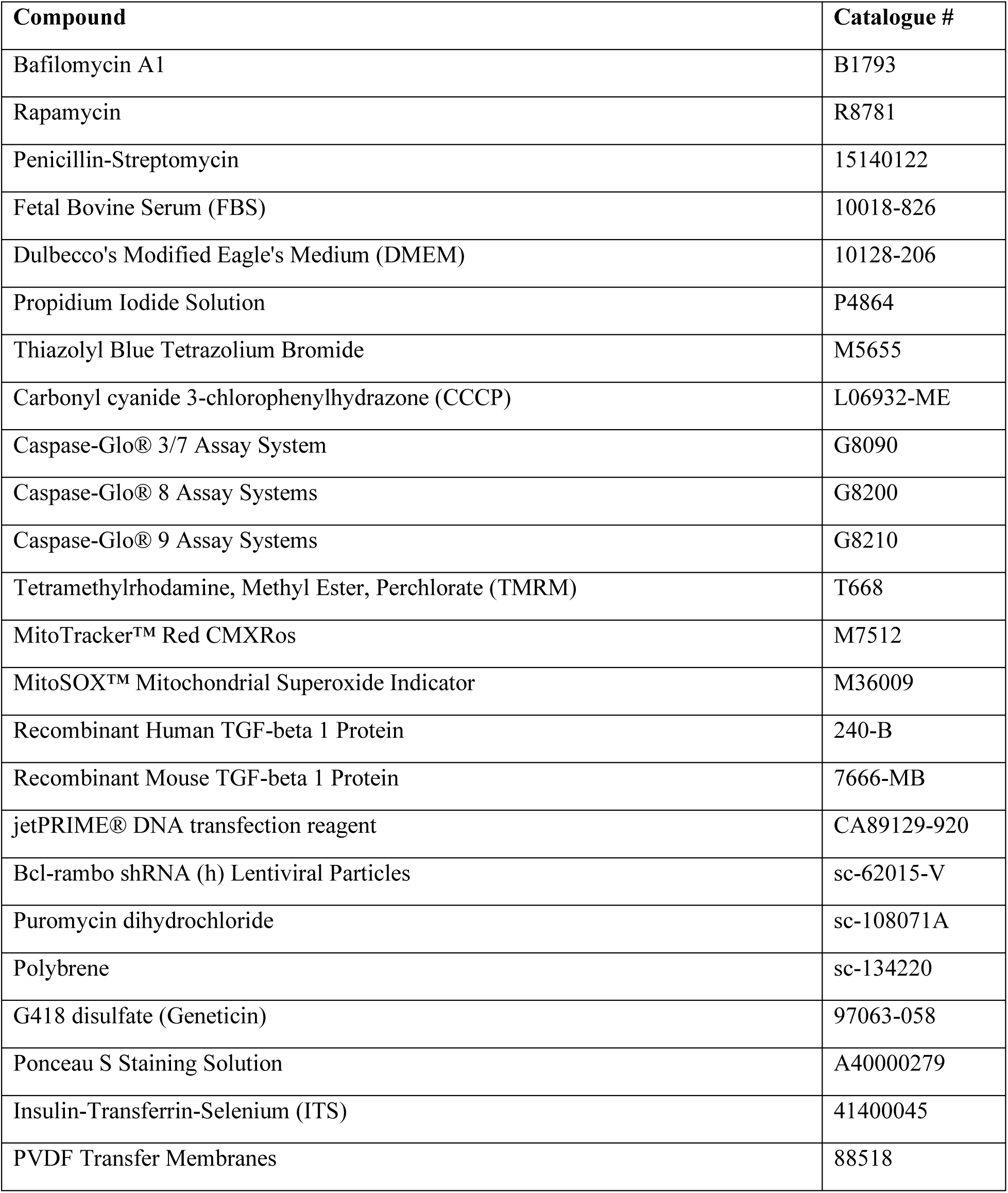
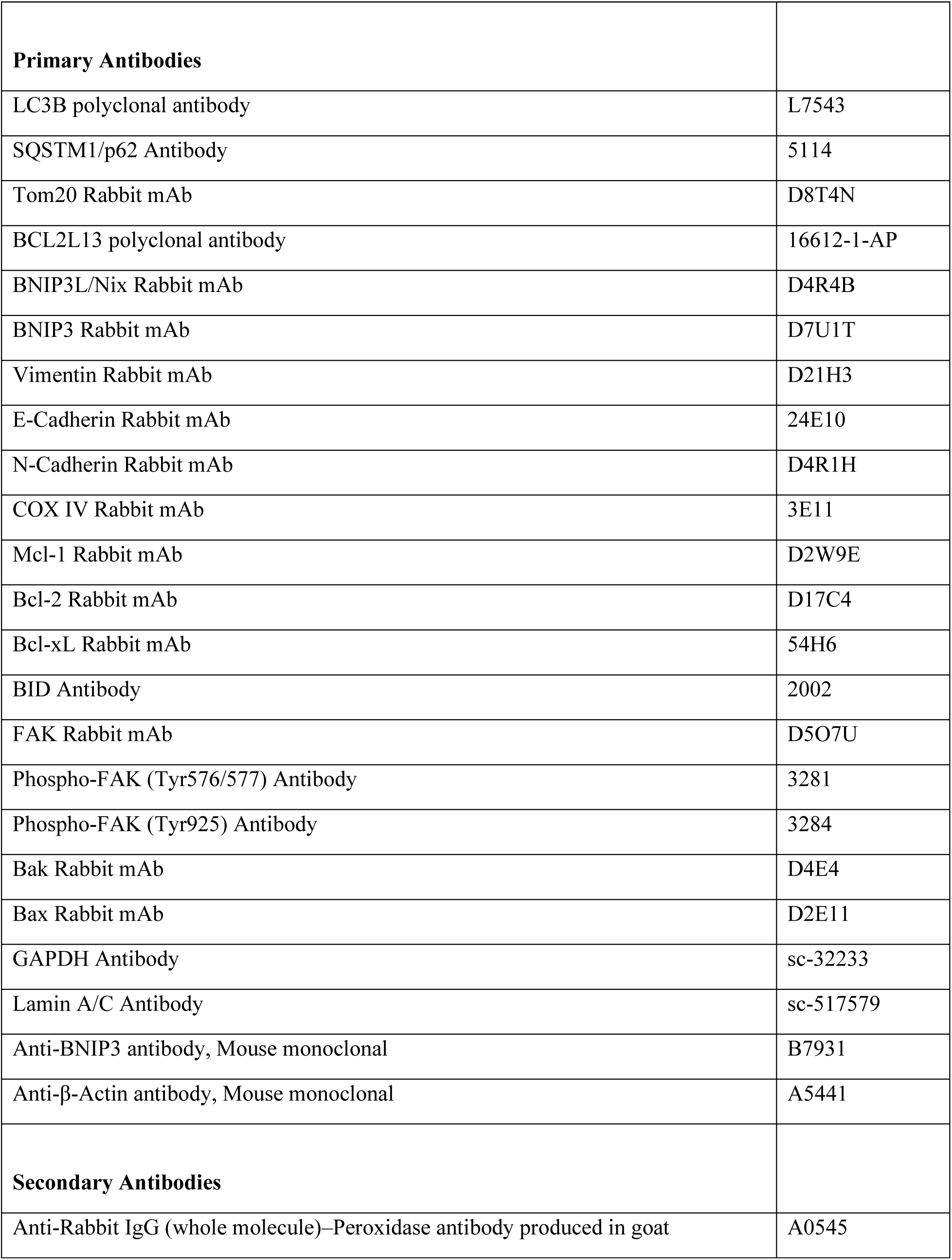
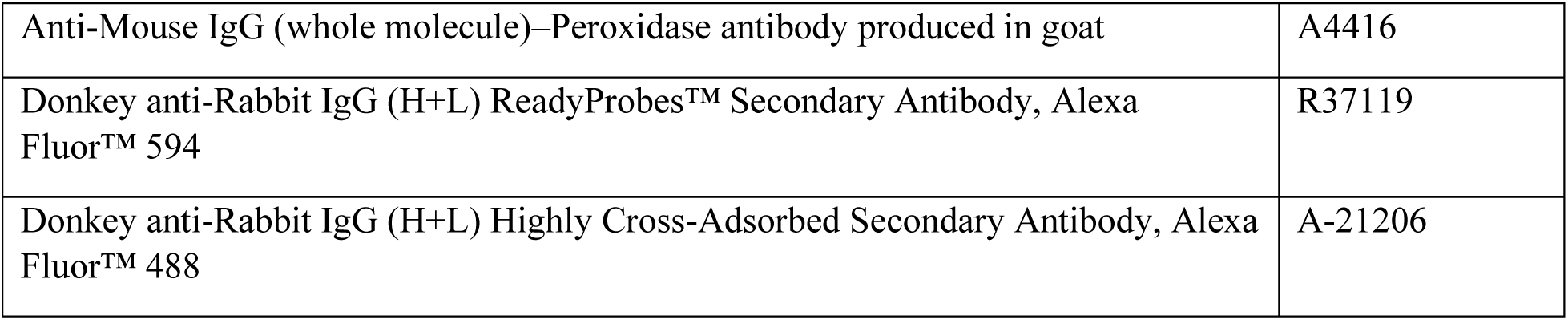
Chemicals, compounds, kits, and antibodies.

| **Compound** | **Catalogue #** |
| --- | --- |
| Bafilomycin A1 | B1793 |
| Rapamycin | R8781 |
| Penicillin-Streptomycin | 15140122 |
| Fetal Bovine Serum (FBS) | 10018-826 |
| Dulbecco's Modified Eagle's Medium (DMEM) | 10128-206 |
| Propidium Iodide Solution | P4864 |
| Thiazolyl Blue Tetrazolium Bromide | M5655 |
| Carbonyl cyanide 3-chlorophenylhydrazone (CCCP) | L06932-ME |
| Caspase-Glo® 3/7 Assay System | G8090 |
| Caspase-Glo® 8 Assay Systems | G8200 |
| Caspase-Glo® 9 Assay Systems | G8210 |
| Tetramethylrhodamine, Methyl Ester, Perchlorate (TMRM) | T668 |
| MitoTracker™ Red CMXRos | M7512 |
| MitoSOX™ Mitochondrial Superoxide Indicator | M36009 |
| Recombinant Human TGF-beta 1 Protein | 240-B |
| Recombinant Mouse TGF-beta 1 Protein | 7666-MB |
| jetPRIME® DNA transfection reagent | CA89129-920 |
| Bcl-rambo shRNA (h) Lentiviral Particles | sc-62015-V |
| Puromycin dihydrochloride | sc-108071A |
| Polybrene | sc-134220 |
| G418 disulfate (Geneticin) | 97063-058 |
| Ponceau S Staining Solution | A40000279 |
| Insulin-Transferrin-Selenium (ITS) | 41400045 |
| PVDF Transfer Membranes | 88518 |
| **Primary Antibodies** |  |
| LC3B polyclonal antibody | L7543 |
| SQSTM1/p62 Antibody | 5114 |
| Tom20 Rabbit mAb | D8T4N |
| BCL2L13 polyclonal antibody | 16612-1-AP |
| BNIP3L/Nix Rabbit mAb | D4R4B |
| BNIP3 Rabbit mAb | D7U1T |
| Vimentin Rabbit mAb | D21H3 |
| E-Cadherin Rabbit mAb | 24E10 |
| N-Cadherin Rabbit mAb | D4R1H |
| COX IV Rabbit mAb | 3E11 |
| Mcl-1 Rabbit mAb | D2W9E |
| Bcl-2 Rabbit mAb | D17C4 |
| Bcl-xL Rabbit mAb | 54H6 |
| BID Antibody | 2002 |
| FAK Rabbit mAb | D5O7U |
| Phospho-FAK (Tyr576/577) Antibody | 3281 |
| Phospho-FAK (Tyr925) Antibody | 3284 |
| Bak Rabbit mAb | D4E4 |
| Bax Rabbit mAb | D2E11 |
| GAPDH Antibody | sc-32233 |
| Lamin A/C Antibody | sc-517579 |
| Anti-BNIP3 antibody, Mouse monoclonal | B7931 |
| Anti-β-Actin antibody, Mouse monoclonal | A5441 |
| **Secondary Antibodies** |  |
| Anti-Rabbit IgG (whole molecule)–Peroxidase antibody produced in goat | A0545 |
| Anti-Mouse IgG (whole molecule)–Peroxidase antibody produced in goat | A4416 |
| Donkey anti-Rabbit IgG (H+L) ReadyProbes™ Secondary Antibody, Alexa Fluor™ 594 | R37119 |
| Donkey anti-Rabbit IgG (H+L) Highly Cross-Adsorbed Secondary Antibody, Alexa Fluor™ 488 | A-21206 |

**Supplementary Table 2.** Antibody dilutions for Immunoblotting and Immunocytochemistry.

| **Primary Antibodies** | **Dilution** |
| --- | --- |
| LC3B polyclonal antibody | 1/2000 |
| SQSTM1/p62 Antibody | 1/1000 |
| Tom20 Rabbit mAb | 1/1000 |
| BCL2L13 polyclonal antibody | 1/2500 |
| BNIP3L/Nix Rabbit mAb | 1/1000 |
| BNIP3 Rabbit mAb | 1/1000 |
| Vimentin Rabbit mAb | 1/1000 |
| E-Cadherin Rabbit mAb | 1/1000 |
| N-Cadherin Rabbit mAb | 1/1000 |
| COX IV Rabbit mAb | 1/1000 |
| Mcl-1 Rabbit mAb | 1/1000 |
| Bcl-2 Rabbit mAb | 1/1000 |
| Bcl-xL Rabbit mAb | 1/1000 |
| BID Antibody | 1/1000 |
| FAK Rabbit mAb | 1/1000 |
| Phospho-FAK (Tyr576/577) Antibody | 1/1000 |
| Phospho-FAK (Tyr925) Antibody | 1/1000 |
| Bak Rabbit mAb | 1/1000 |
| Bax Rabbit mAb | 1/1000 |
| GAPDH Antibody | 1/500 |
| Lamin A/C Antibody | 1/500 |
| BNIP3 antibody, Mouse monoclonal | 1/1000 |
| β-Actin antibody, Mouse monoclonal | 1/10,000 |
| **Secondary Antibodies** |  |
| Anti-Rabbit IgG (whole molecule)–Peroxidase antibody produced in goat | 1/2000 |
| Anti-Mouse IgG (whole molecule)–Peroxidase antibody produced in goat | 1/2000 |
| Donkey anti-Rabbit IgG (H+L) ReadyProbes™ Secondary Antibody, Alexa Fluor™ 594 | 1/2000 |
| Donkey anti-Rabbit IgG (H+L) Highly Cross-Adsorbed Secondary Antibody, Alexa Fluor™ 488 | 1/2000 |

